# Inducible Depletion of Calpain-2 Mitigates Abdominal Aortic Aneurysm in Mice

**DOI:** 10.1101/2020.04.15.043687

**Authors:** Latha Muniappan, Michihiro Okuyama, Aida Javidan, Devi Thiagarajan, Weihua Jiang, Jessica J. Moorleghen, Lihua Yang, Anju Balakrishnan, Deborah A. Howatt, Haruhito A. Uchida, Takaomi C. Saido, Venkateswaran Subramanian

## Abstract

**BACKGROUND:** Cytoskeletal structural proteins maintain cell structural integrity by bridging extracellular matrix (ECM) with contractile filaments. During AAA development, (i) aortic medial degeneration is associated with loss of smooth muscle cell (SMC) integrity, and (ii) fibrogenic mesenchymal cells (FMSCs) mediates ECM remodeling. Calpains cleave cytoskeletal proteins that maintain cell structural integrity. Pharmacological inhibition of calpains exert beneficial effects on Angiotensin II (AngII)-induced AAAs in low density receptor deficient (LDLR-/-) mice.

**OBJECTIVES:** To evaluate the functional contribution of FMSCs-derived calpain-2 on (i) cytoskeletal structural protein and ECM alterations, and (ii) AAA progression.

**METHODS:** Calpain-2 protein, and cytoskeletal protein (e.g. filamin or talin) fragmentation in human and mice AAA tissues were assessed by immunohistochemical and western blot analyses. LDLR-/- mice that were either inducible-whole body or FMSC-specific calpain-2 deficient were fed a fat-enriched diet and infused with AngII for 4 weeks. The association of cytoskeletal protein to ECM was evaluated using aortic SMCs, in vitro. In addition, the effect of calpain-2 deficiency on the stability of established AAA was examined.

**RESULTS:** Calpain-2 protein, and filamin/talin fragmentation are significantly elevated in AAAs. Ubiquitous or FMSC-specific depletion of calpain-2 suppressed AngII-induced AAAs, filamin/talin fragmentation and promoted ECM protein, collagen. Calpain-2 silencing in SMCs reduced AngII-induced filamin/talin fragmentation. In addition, silencing of filamin or talin in SMCs significantly reduced collagen protein. Furthermore, calpain-2 deficiency suppressed established AAA rupture.

**CONCLUSION:** Calpain-2 activation promotes cytoskeletal structural protein fragmentation and ECM degradation of experimental AAA aortas. Treatment with calpain-2 specific inhibitor may facilitate the clinical management of AAA.

## INTRODUCTION

Abdominal aortic aneurysms (AAAs) are permanent dilations of the abdominal aorta with greater than 80% mortality after rupture. (1) Lack of sufficient mechanistic knowledge and availability of specific therapeutic agents to stabilize and /or prevent aortic rupture, highlight the need to gain mechanistic insights into AAA development to improve the long-term prognosis. Loss of smooth muscle cell (SMC) integrity is associated with aortic medial degeneration during AAA formation. (2-4) Cytoskeletal structural proteins maintain cell structural integrity by bridging the extracellular matrix (ECM) with SMC contractile filaments. (5-8) Activated mesenchymal cells which includes adventitial fibroblasts, myofibroblasts and a subpopulation of synthetic medial SMCs play a potent role on ECM remodeling. These heterogenous lineage cells secrete ECM protein, type I collagen which is composed of α1 and α2 peptides, and encoded by Col1α1 and Col1α2.(9, 10) Currently, it is not clear whether cytoskeletal structural proteins play a role in vessel wall integrity during AAA development, and the identity of intracellular proteases especially of the fibrogenic mesenchymal cells origin that targets cytoskeletal structural proteins remain undefined.

Calpains, calcium-dependent cysteine proteases, tightly regulate target protein substrates through limited proteolysis.(11) The two major isoforms, calpain-1 and −2, are expressed ubiquitously; whereas the other isoforms (e.g. −3, −9) are tissue-specific.(11) Calpains are the only known intracellular proteases that target an array of cytoskeletal and membrane proteins.(11-14) Earlier, we demonstrated that pharmacological inhibition of calpain-1 and −2 reduced angiotensin II (AngII)-induced AAAs in mice.(15) In addition, calpain-2 compensates for the loss of calpain-1 and promoted AAA formation in calpain-1 deficient mice.(16) Furthermore, using leukocyte-specific calpain-2 deficient mice, we demonstrated that leukocyte-derived calpain-2 had no influence on AngII-induced AAA development in mice.(17)

In this study, using human AAA tissue samples, we examined calpain-2 protein and activity and fragmentation of cytoskeletal structural proteins, filamin A and talin. In addition, to elucidate the functional role of calpain-2 in AngII-induced AAA development and cytoskeletal protein fragmentation, we generated an inducible whole body or mesenchymal-specific calpain-2 deficient mice in an LDL receptor deficient background. These studies demonstrated that (i) calpain-2 protein and activity, and cytoskeletal structural protein fragmentation were increased in human and experimental AAAs, (ii) induced depletion of calpain-2 strongly suppressed AAA development and cytoskeletal protein fragmentation in mice, (iii) mesenchymal derived calpain-2 deficiency strongly suppressed AAA development in mice. Furthermore, siRNA mediated silencing of calpain-2 in aortic SMCs prevented AngII-induced filamin and talin fragmentation. In addition, to elucidate the functional role of cytoskeletal structural proteins, filamin A and talin, on ECM protein stability, we examined the effect of filamin A or talin silencing in aortic SMCs. These studies demonstrated that silencing of either filamin A or talin significantly reduced the abundance of collagen protein in aortic SMCs. Furthermore, inducible depletion of calpain-2 also prevented rupture of established AAAs with improved survival rate.

## RESULTS

### Calpain-2 Protein and Activity are increased in Human and Experimental AAAs

In the present study, immunohistochemical staining of human AAA sections showed a strong presence of calpain-2 protein compared to control abdominal aortic sections (Figure 1A-L). The immunohistochemical staining showed a strong distribution of calpain-2 in fibroblast rich adventitia, surrounding perivascular adipose tissue, and in infiltrated macrophages, with a weak staining in the intima and media of human AAA. Similarly, immunohistochemical staining on AngII-induced AAAs showed a strong distribution of calpain-2 in infiltrated macrophages, surrounding perivascular adipose tissue, and fibroblast rich adventitia, with a weak staining in the intima and media (Figure 1M-O). Furthermore, Western blot analyses showed a significant increase in calpain-2 protein in both human (Figure 1P,Q) and AngII-induced AAAs compared to control tissues (Figure 1S,T). In addition, measurement of calpain activity using a fluorescent substrate of calpain, Ac-LLY-AFC, showed a significant increase in calpain activity in both human (Figure 1R) and experimental AAA tissues (Figure 1U) compared to controls.

**Figure 1.**
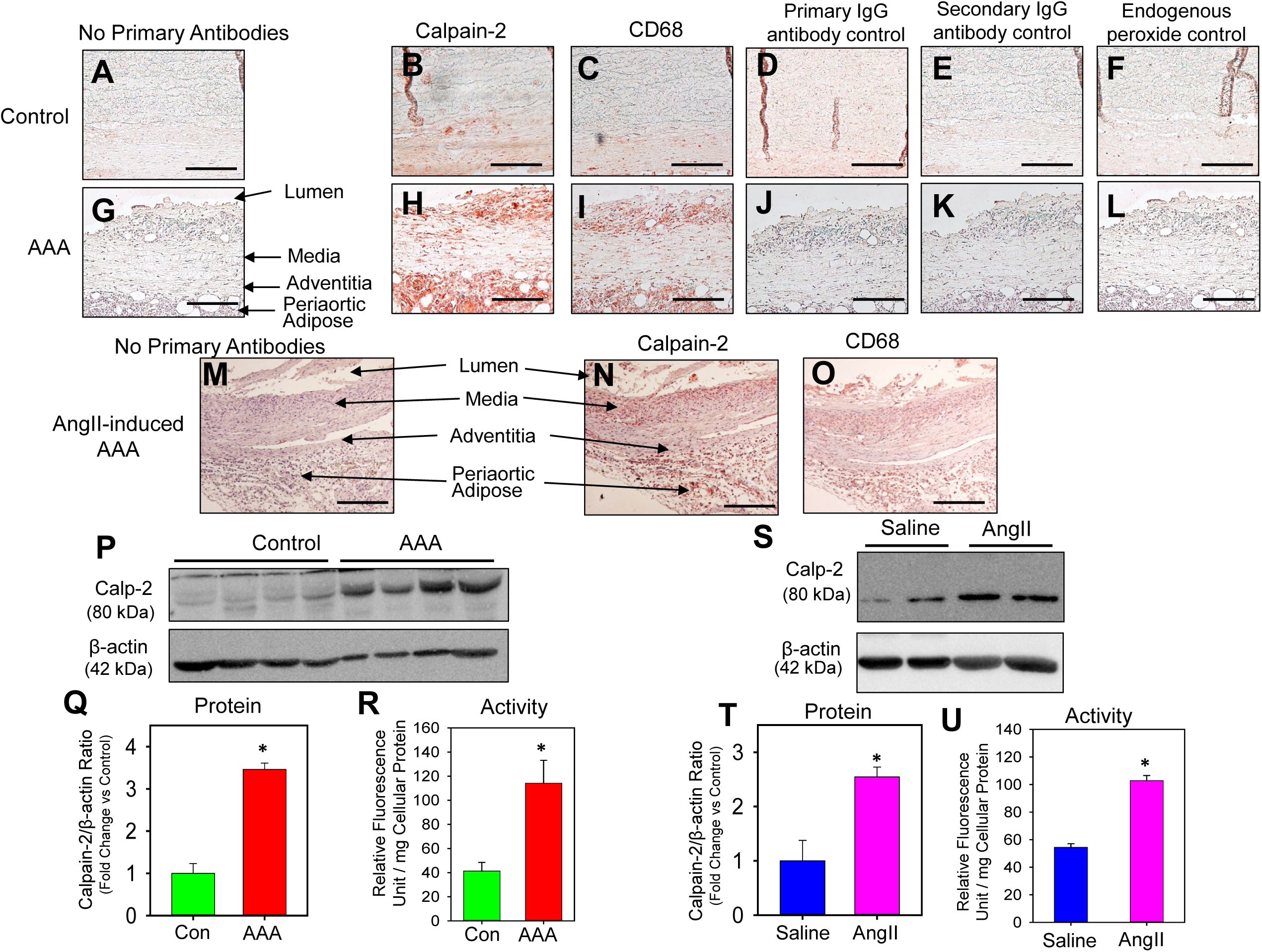
Calpain-2 protein and activity are increased in human and experimental AAAs. Serial cross-sections from human AAA and control abdominal aortic tissue sections (**A, G**) were immunostained with anti-calpain-2 (**B, H**) and CD68 (**C, I**). Red color indicates positive immunostaining. Representative control and AAA tissue sections stained with rat and rabbit primary IgG iso-type control antibodies (**D, J**), secondary antibodies (Rabbit anti-rat or Goat anti-rabbit-**E,K**) and endogenous peroxide (no antibodies - **F,L**) control. Serial cross-sections from AngII-induced AAA tissue sections were immunostained with anti-calpain-2 (**M,N**) and CD68 (**O**). Nuclei were counterstained with hematoxylin. Scale bars correspond to 50 μm (200x magnification). Calpain-2 protein detection in human control abdominal aorta and AAA tissues (n=4) (**P,Q**). Calpain activity (**R**) in human control and AAA tissues (n=6). Calpain-2 and β-actin protein in abdominal aortas from saline and AngII-infused LDL receptor-/- mice (n=6) (**S,T**). Calpain activity (**U**) in abdominal aortas from saline and AngII-infused LDL receptor-/- mice (n=4). * denotes P<0.05 when comparing control vs AAA or saline vs AngII infusion (Student’s *t* test).

### Generation of Inducible Calpain-2 Deficient Mice

Since calpain-2 deficiency in mice is embryonically lethal,(18) calpain-2 deficient LDL receptor -/- mice were generated using calpain-2 floxed (f/f) and Cre transgenic mice expressing a tamoxifen-inducible Cre recombinase under the control of the ubiquitously expressed chicken β-actin promoter (ERT2 Cre+/0).(19-21) Both calpain-2 f/f and β-actin Cre transgenic mice were bred to an LDL receptor -/- background. Female calpain-2 f/f mice were bred with male β-actin Cre transgenic mice to yield offspring homologous for the floxed allele and hemizygous for the Cre transgene (Cre+/0). Littermates that were homozygous for the floxed calpain-2 gene, but without the Cre transgene (Cre0/0), were used as control mice. Calpain-2 f/f and Cre genotypes were confirmed by PCR (Supplemental Figure S1). To induce Cre recombinase activity, mice were injected with tamoxifen (25 mg/kg body weight) intraperitoneally for 5 consecutive days. Western blot analyses of aortic tissue protein showed complete depletion of capain-2 protein, without influencing calpain-1, in aortas from Cre+/0 mice (Figure 2A).

**Figure 2.**
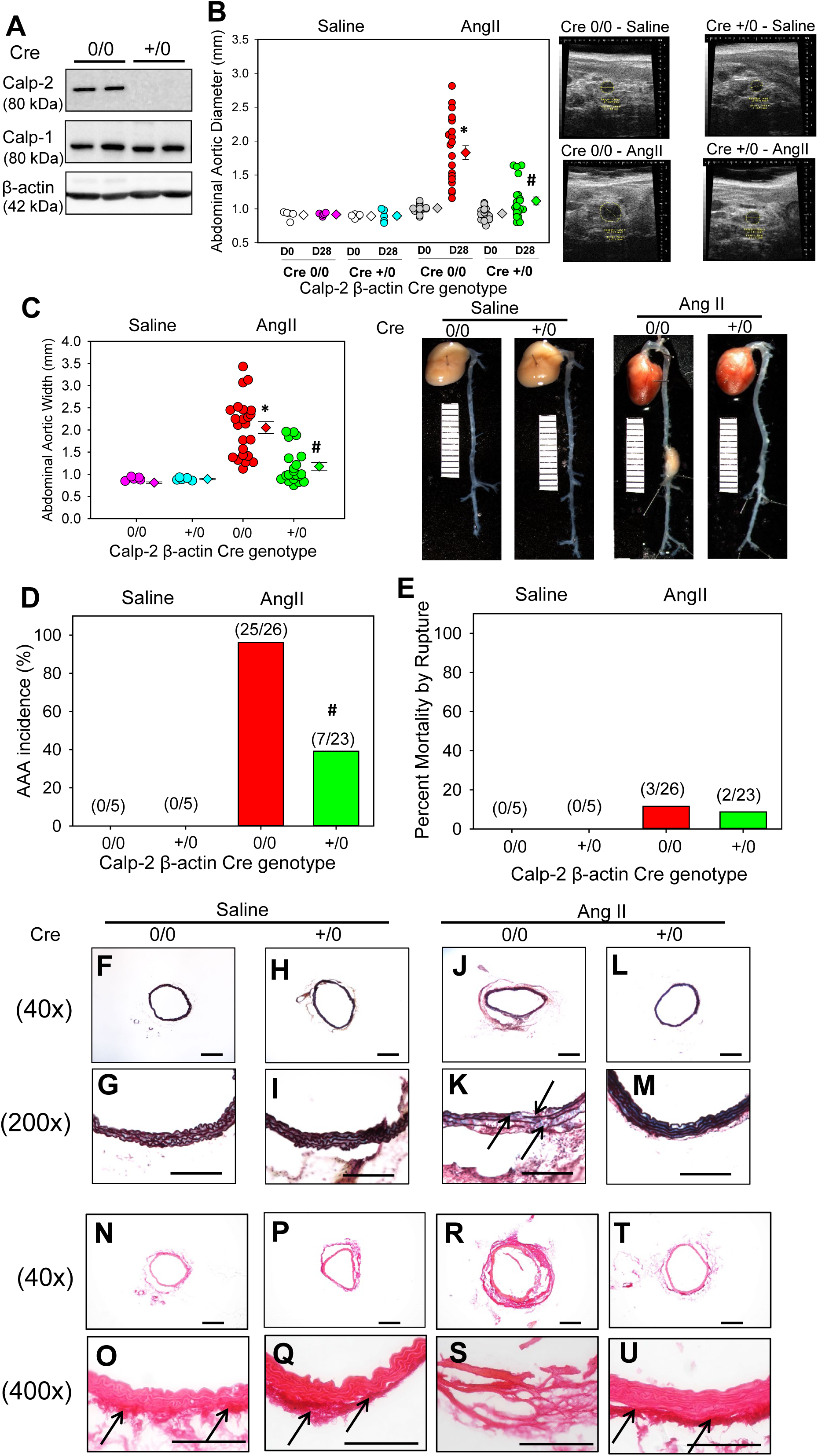
Calpain-2 deficiency significantly reduced AngII-induced AAAs. **A**. Calpain-2 and β-actin protein were detected in aortic tissue lysates from calpain-2 f/f β-actin (ERT2) Cre0/0 and +/0 mice. **B**. Ultrasonic measurements of abdominal aortic luminal diameters were measured on day 0 and after 28 days of saline or AngII infusion (Saline n=5; AngII n=23-26). Measurements of maximal external width of abdominal aortas (Saline n=5; AngII n=23-26). Pink (saline - Cre 0/0), teal (saline - Cre +/0), red (AngII - Cre 0/0) and green (AngII - Cre +/0) represent individual mice, diamonds represent means, and bars are SEMs. * denotes P<0.05 when comparing saline vs AngII infusion; # denotes P<0.05 when comparing Cre 0/0 vs Cre +/0 mice (Two-way ANOVA with Holm-Sidak post hoc analysis). The incidence of AAA (>50% increase in aortic width) in AngII-infused calpain-2 f/f mice that were either β-actin Cre 0/0 (red bar) or Cre +/0 (green bar). Statistical analyses were performed by Fisher Exact test (# denotes P<0.001 when comparing Cre 0/0 vs Cre +/0 mice).**E**. Mortality due to AAA rupture in AngII-infused calpain-2 f/f mice that were either β-actin Cre 0/0 or +/0. Red (Cre0/0) and green bar (Cre+/0). Representative suprarenal aortic tissue-sections from saline (**F-I;N-Q**) and AngII (**J-M;R-U**) administered calpain-2 f/f mice that were β-actin Cre 0/0 and +/0 stained with Movat’s Pentachrome or Picrosirius red. Arrows indicates medial breaks. Scale bars correspond to 50 μm. F,H,J, L, N,P,R and T = 40x; G,I,K, M, O,Q,S and U = 200x.

### Inducible Depletion of Calpain-2 Reduced AngII-induced AAA in Mice

To determine the role of calpain-2 in AngII-induced AAA, calpain-2 f/f mice that are Cre0/0 or +/0 were fed a saturated fat-enriched diet and infused with saline or AngII for 4 weeks. Inducible calpain-2 deficiency had no effect on body weight, total plasma cholesterol concentrations or systolic blood pressure (Table 1). Calpain-2 deficiency significantly reduced AngII-induced aortic luminal dilation (Figure 2B) as measured by ultrasound (Diameter: Saline - Cre0/0: 0.80 ± 0.02 *vs* Cre+/0: 0.83 ± 0.01 P=NS, AngII - Cre0/0: 1.83 ± 0.10 *vs* Cre+/0: 1.10 ± 0.06, P<0.001; Two-way ANOVA). In addition, calpain-2 deficiency significantly reduced AngII-induced AAA formation (Figure 2C) as measured by external aortic width expansion (Mean width of abdominal aorta – Saline - Cre0/0: 0.81 ± 0.02 mm *vs* Cre+/0: 0.89 ± 0.01 mm, P=NS; AngII - Cre0/0: 2.05 ± 0.13 mm *vs* Cre+/0: 1.17 ± 0.09 mm, P<0.001; Two-way ANOVA). Furthermore, calpain-2 deficiency also significantly reduced AngII-induced AAA incidence (Figure 2D; defined as a >50% increase in suprarenal aorta width; Cre0/0: 91% *vs* Cre+/0: 30%, P<0.001; Fisher Exact Test) but had no significant effect on aortic rupture (Figure 2E; Cre0/0: 11% *vs* Cre+/0: 8%).

**Table 1.** Effects of calpain-2 deficiency in male LDL receptor -/- mice infused with saline or AngII.

| Groups | Cre 0/0 |  | Cre +/-0 |  |
| --- | --- | --- | --- | --- |
|  | Saline | AngII | Saline | AngII |
| N | 5 | 26 | 5 | 23 |
| Body Weight (g) | 31.3 ± 1.1 | 30.4 ± 0.7 | 29.6 ± 0.9 | 29.4 ± 0.5 |
| Plasma Cholesterol (mg/dL) | 1458 ± 92 | 1597 ± 43 | 1291 ± 60 | 1541 ± 63 |
| Systolic BP Pre-Infusion (mmHg) | 139 ± 6 | 141 ± 4 | 140 ± 3 | 146 ± 4 |
| Systolic BP Post-Infusion (mmHg) | 141 ± 4 | 187 ± 5* | 139 ± 4 | 190 ± 3* |

Values are represented as means ± SEMs. Body weights and plasma cholesterol concentrations were determined at termination. Systolic blood pressure was measured prior to (week 0) and during AngII infusion (week 4). Two way repeated measures ANOVA was used to analyze systolic blood pressures. \* Denotes $P < 0.05$ systolic BP post-infusion vs pre-infusion, by two-way repeated measures ANOVA. # Denotes $P < 0.05$ AngII vs saline by two-way ANOVA with Holm-Sidak post hoc analysis.

Histological staining of abdominal aortas using Movat’s Pentachrome staining (Figure 2F-M) revealed occurrence of focal elastin layer disruption (Figure 2J, K) in the Cre 0/0 group infused with AngII. However, the abdominal aortas from calpain-2 deficient mice (Cre+/0) showed preserved the medial elastin layer upon AngII infusion (Figure 2L,M). Similarly, picrosirius red staining revealed that abdominal aortas from calpain-2 deficient mice had more adventitial collagen compared to Cre0/0 group (Figure 2 P,Q,T,U). In addition, calpain-2 deficiency had no effect on AngII-induced atherosclerotic lesion areas in aortic arches (Cre0/0-17.3 ± 1.7 % *vs* Cre+/0 – 17.7 ± 2.0 %; Supplemental Figure S2) and thorax (Cre0/0-4.7 ± 0.6 % *vs* Cre+/0 – 4.8 ± 0.6 %; Supplemental Figure S2).

### Calpain-2 Deficiency in Adipocytes had no Influence on AngII-induced AAA Formation in Mice

In mice, evidences suggest that obesity-accelerated perivascular inflammation play a critical role in the acceleration of AngII-induced AAA formation.(22) Further, human AAA tissue characterization studies showed an accelerated inflammation in the surrounding periaortic adipose tissue. Based on the strong calpain-2 positive staining observed in both human and AngII-induced AAAs, next we sought to determine the contribution of adipocyte-derived calpain-2 in the development of AngII-induced AAAs. Adipocyte-specific calpain-2 deficient LDL receptor -/- mice were generated using calpain-2 floxed (f/f) and Cre transgenic mice expressing Cre recombinase under the control of the adipocyte-specific adiponectin (Adipoq) promoter.Both calpain-2 f/f and Adipoq Cre transgenic mice were bred to an LDL receptor -/- background. Female calpain-2 f/f mice were bred with male Adipoq Cre transgenic mice to yield offspring homologous for the floxed allele and hemizygous for the Cre transgene (Cre+/0). Littermates that were homozygous for the floxed calpain-2 gene, but without the Cre transgene (Cre0/0), were used as control mice. Calpain-2 f/f and Cre genotypes were confirmed by PCR (Supplemental Figure S3). Western blot analyses of various fat pads (retroperitoneal, brown adipose, periaortic adipose) and aorta showed a strong reduction of calpain-2 in adipose tissue with intact calpain-2 in the aorta (Figure 3A).

**Figure 3.**
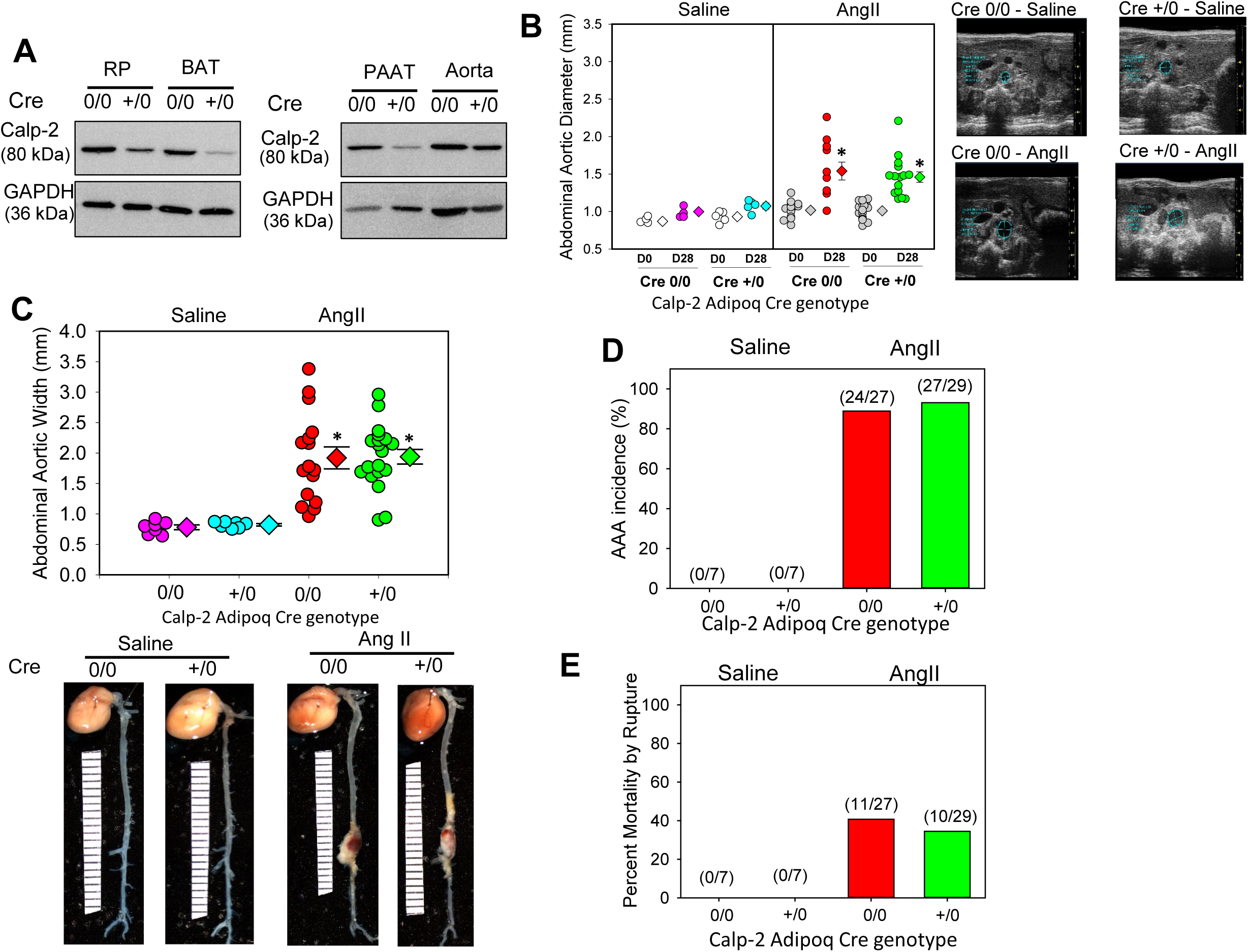
Calpain-2 deficiency in adipocytes had no influence on AngII-induced AAAs. **A**. Calpain-2 and β-actin protein were detected in various fat pads (retroperitoneal, brown adipose, periaortic adipose) and aortic tissue lysates from calpain-2 f/f Adipoq Cre 0/0 and +/0 mice. **B**. Ultrasonic measurements of abdominal aortic luminal diameters were measured on day 0 and after 28 days of saline or AngII infusion (Saline n=5; AngII n=23-27). **C**. Measurements of maximal external width of abdominal aortas (Saline n=6-7; AngII n=27-29). Pink (saline - Cre 0/0), teal (saline - Cre +/0), red (AngII - Cre 0/0) and green (AngII - Cre +/0) represent individual mice, diamonds represent means, and bars are SEMs. * denotes P<0.05 when comparing saline vs AngII infusion; # denotes P<0.05 when comparing Cre 0/0 vs Cre +/0 mice (Two-way ANOVA with Holm-Sidak post hoc analysis). **D**. The incidence of AAA (>50% increase in aortic width) in AngII-infused calpain-2 f/f mice that were either Adipoq Cre 0/0 (red bar) or Cre +/0 (green bar). Statistical analyses were performed by Fisher Exact test (# denotes P<0.001 when comparing Cre 0/0 vs Cre +/0 mice). **E**. Mortality due to AAA rupture in AngII-infused calpain-2 f/f mice that were either Adipoq Cre 0/0 or +/0. Red (Cre0/0) and green bar (Cre+/0).

To determine the role of adipocyte-calpain-2 in AngII-induced AAA, calpain-2 f/f mice that are Adipoq Cre0/0 or +/0 were fed a saturated fat-enriched diet and infused with saline or AngII for 4 weeks. Adipocyte-specific calpain-2 deficiency had no effect on body weight or total plasma cholesterol concentrations (Table 2). Calpain-2 deficiency in adipocytes had no effect on AngII-induced aortic luminal dilation (Figure 3B; Diameter: Saline - Cre0/0: 0.87 ± 0.01 *vs* Cre+/0: 0.99 ± 0.02 P=NS, AngII - Cre0/0: 1.54 ± 0.12 *vs* Cre+/0: 1.46 ± 0.07, P=NS; Two-way ANOVA), and external aortic width expansion (Figure 3C; Mean width of abdominal aorta – Saline - Cre0/0: 0.78 ± 0.04 mm *vs* Cre+/0: 0.82 ± 0.02 mm, P=NS; AngII - Cre0/0: 1.92 ± 0.18 mm *vs* Cre+/0: 1.94 ± 0.12 mm, P=NS; Two-way ANOVA). Furthermore, calpain-2 deficiency in adipocytes showed no difference on AngII-induced AAA incidence and aortic rupture (Figure 3 D and E). In addition, calpain-2 deficiency in adipocytes had no effect on AngII-induced atherosclerotic lesion areas in aortic arches (Cre0/0-12.0 ± 1.0 % *vs* Cre+/0 – 10.9 ± 1.10 %; Online Figure 4 in the online-only Data Supplement).

**Table 2.** Effects of adipocyte-calpain-2 deficiency in male LDL receptor -/- mice infused with saline or AngII.

| **Groups** | **Cre 0/0** |  | **Cre +/-0** |  |
| --- | --- | --- | --- | --- |
| **Infusion** | **Saline** | **AngII** | **Saline** | **AngII** |
| N | 6 | 27 | 7 | 29 |
| Body Weight (g) | 28.4 ± 0.9 | 24.4 ± 0.4 | 29.1 ± 1.0 | 24.2 ± 0.5 |
| Plasma Cholesterol (mg/dL) | 1418 ± 62 | 1305 ± 45 | 1384 ± 60 | 1265 ± 58 |

Values are represented as means ± SEMs. Body weights and plasma cholesterol concentrations were determined at termination. \* Denotes $P < 0.05$ AngII vs saline by two-way ANOVA with Holm-Sidak post hoc analysis.

**Figure 4.**
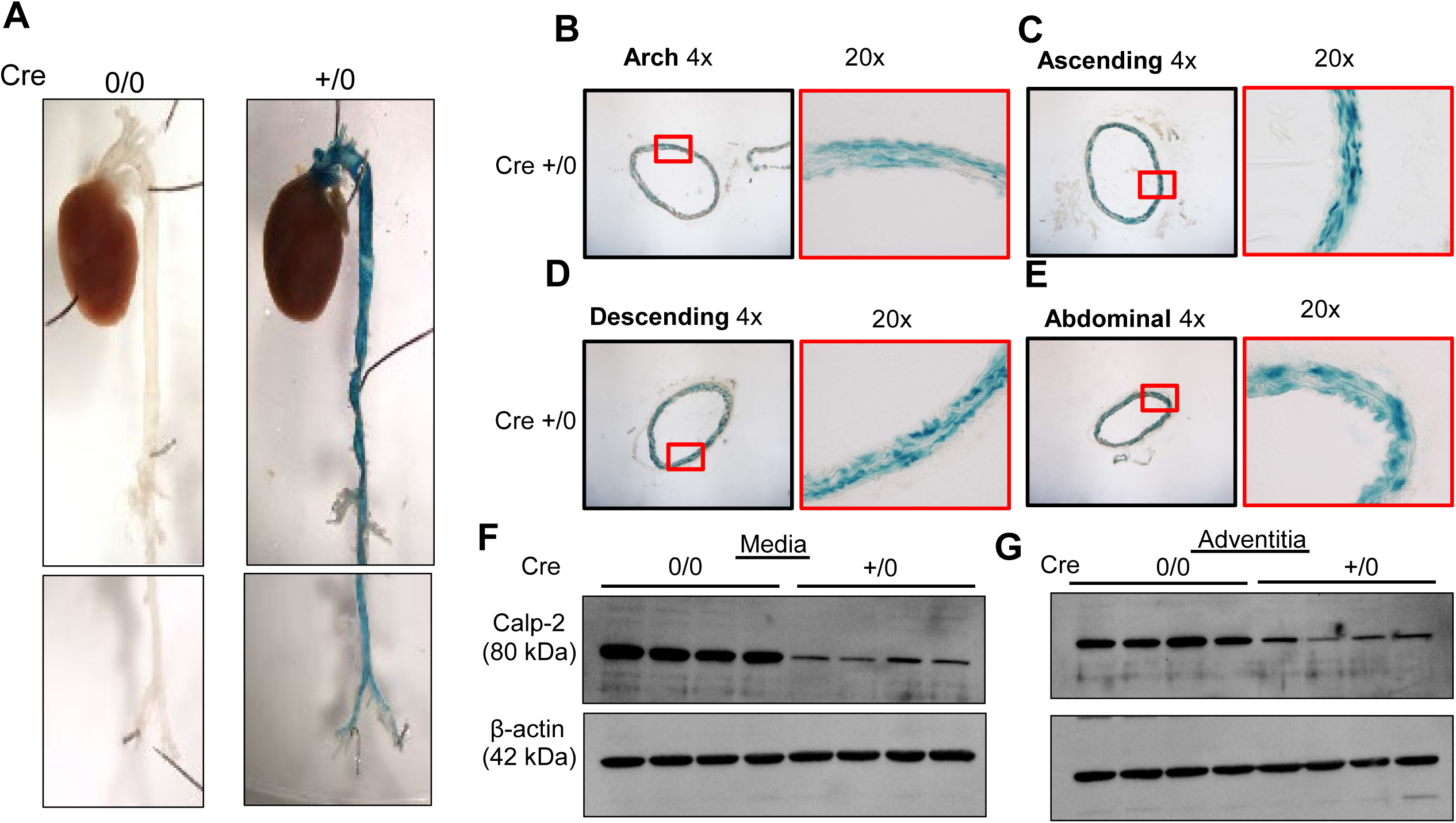
Distribution of Col1a2 positive mesenchymal cells in the aorta. **A**. Representative ventral views of β-galactosidase (β-gal) activity in aortas from Col1a2 Cre positive (Cre+/0) and control (Cre0/0) mice. Blue color is positive staining for distribution of Cre excision. Cross-sections of arch (**B**), ascending (**C**), descending (**D**) and abdominal (**E**) aortas from Col1a2 Cre mice stained with X-gal. Calpain-2 and β-actin protein were detected in aortic medial (**F**) and adventitial (**G**) tissue lysates from calpain-2 f/f Col1a2 Cre 0/0 and +/0 mice.

### Inducible Depletion of Calpain-2 in Col1a2 Positive Mesenchymal Cells Reduced AngII-induced AAA in Mice

Previous studies demonstrated that tamoxifen mediated induction of Col1a2 Cre-recombinase (Col1a2-ERT2) resulted in the activation of Cre recombinase in the fibrogenic mesenchymal cells (aortic vascular SMCs and fibroblasts).(10, 23, 24) Since many cell-specific promoters have promiscuous expression that led to broader expression than defined by nomenclature, first, we determined the distribution of Col1a2-ERT2 Cre recombinase by breeding these transgenic mice to ROSA26^LacZ^ reporter mice. Whole aortic tissue showed presence of β-galactosidase (β-gal) activity in Col1a2-ERT2 Cre+/0 mice (Figure 4A), whereas no positive activity was detected in non-transgenic littermates (Cre0/0). Furthermore, to determine the distribution of Col1a2-ERT2 Cre-lacZ in the aorta, we serially sectioned the aorta and found that β-gal is strongly distributed in the SMC rich aortic media and fibroblast rich adventitia throughout the aorta (Figure 4 B-E). In addition to aorta, presence of β-gal in fibroblast and SMC rich organs (e.g. bladder, intestine, lung, skin etc.) and absence of β-gal in non-fibrotic organs such as brain, thymus (Supplemental Figure S5) confirm the specificity of Col1a2-ERT2 Cre.

To develop fibrogenic mesenchymal cell specific-calpain-2 deficient mice, both calpain-2 f/f and Col1a2-ERT2 Cre transgenic mice were bred to an LDL receptor -/- background. Female calpain-2 f/f mice were bred with male Col1a2-ERT2 Cre transgenic mice to yield offspring homologous for the floxed allele and hemizygous for the Cre transgene (Cre+/0). Littermates that were homozygous for the floxed calpain-2 gene, but without the Cre transgene (Cre0/0), were used as control mice. Calpain-2 f/f and Cre genotypes were confirmed by PCR (Supplemental Figure S6). To induce Cre recombinase activity, mice were injected with tamoxifen (25 mg/kg body weight) intraperitoneally for 5 consecutive days. Western blot analyses of SMC-rich aortic medial or fibroblast rich aortic adventitial tissue protein showed a dramatic reduction of capain-2 protein, in Cre+/0 mice (Figure 4 F and G).

To determine the role of mesenchymal cell-derived calpain-2 in AngII-induced AAA, calpain-2 f/f mice that are Cre0/0 or +/0 were fed a saturated fat-enriched diet and infused with saline or AngII for 4 weeks. Inducible calpain-2 deficiency in mesenchymal cells had no effect on body weight, total plasma cholesterol concentrations or systolic blood pressure (Table 3). Calpain-2 deficiency significantly reduced AngII-induced aortic luminal dilation (Figure 5A) as measured by ultrasound (Diameter: Saline - Cre0/0: 1.0 ± 0.02 *vs* Cre+/0: 1.1 ± 0.02 P=NS, AngII - Cre0/0: 2.0 ± 0.15 *vs* Cre+/0: 1.52 ± 0.13, P<0.05; Two-way ANOVA). In addition, calpain-2 deficiency significantly reduced AngII-induced AAA formation (Figure 5B and C) as measured by external aortic width expansion (Mean width of abdominal aorta – Saline - Cre0/0: 0.90 ± 0.03 mm *vs* Cre+/0: 0.83 ± 0.04 mm, P=NS; AngII - Cre0/0: 2.18 ± 0.15 mm *vs* Cre+/0: 1.37 ± 0.13 mm, P<0.05; Two-way ANOVA). Furthermore, calpain-2 deficiency also significantly reduced AngII-induced AAA incidence (Figure 5D; Cre0/0: 98% *vs* Cre+/0: 50%, P<0.001; Fisher Exact Test) but had no significant effect on aortic rupture (Figure 5E; Cre0/0: 39% *vs* Cre+/0: 15%, P=0.09; Fisher Exact Test). In addition, calpain-2 deficiency in fibrogenic mesenchymal cells had no effect on AngII-induced atherosclerotic lesion areas in aortic arches (Cre0/0-11.8 ± 1.6 % *vs* Cre+/0 – 12.8 ± 1.23 %; Supplemental Figure S7).

**Table 3.** Effects of Col1a2 positive mesenchymal cell-specific calpain-2 deficiency in male LDL receptor -/- mice infused with saline or AngII.

| **Groups** | **Cre 0/0** |  | **Cre +/-0** |  |
| --- | --- | --- | --- | --- |
| **Infusion** | **Saline** | **AngII** | **Saline** | **AngII** |
| N | 5 | 23 | 5 | 20 |
| Body Weight (g) | 30.6 ± 1.0 | 25.6 ± 0.5 | 30.4 ± 1.5 | 24.7 ± 0.3 |
| Plasma Cholesterol (mg/dL) | 1042 ± 83 | 884 ± 52 | 866 ± 83 | 842 ± 44 |
| Systolic BP Pre-Infusion (mmHg) | 147 ± 2 | 145 ± 3 | 147 ± 5 | 145 ± 4 |
| Systolic BP Post-Infusion (mmHg) | 148 ± 4 | 185 ± 5* | 158 ± 5 | 187 ± 4* |

Values are represented as means ± SEMs. Body weights and plasma cholesterol concentrations were determined at termination. Systolic blood pressure was measured prior to (week 0) and during AngII infusion (week 4). Two way repeated measures ANOVA was used to analyze systolic blood pressures. \* Denotes $P < 0.05$ systolic BP post-infusion vs pre-infusion, by two-way repeated measures ANOVA. # Denotes $P < 0.05$ AngII vs saline by two-way ANOVA with Holm-Sidak post hoc analysis.

**Figure 5.**
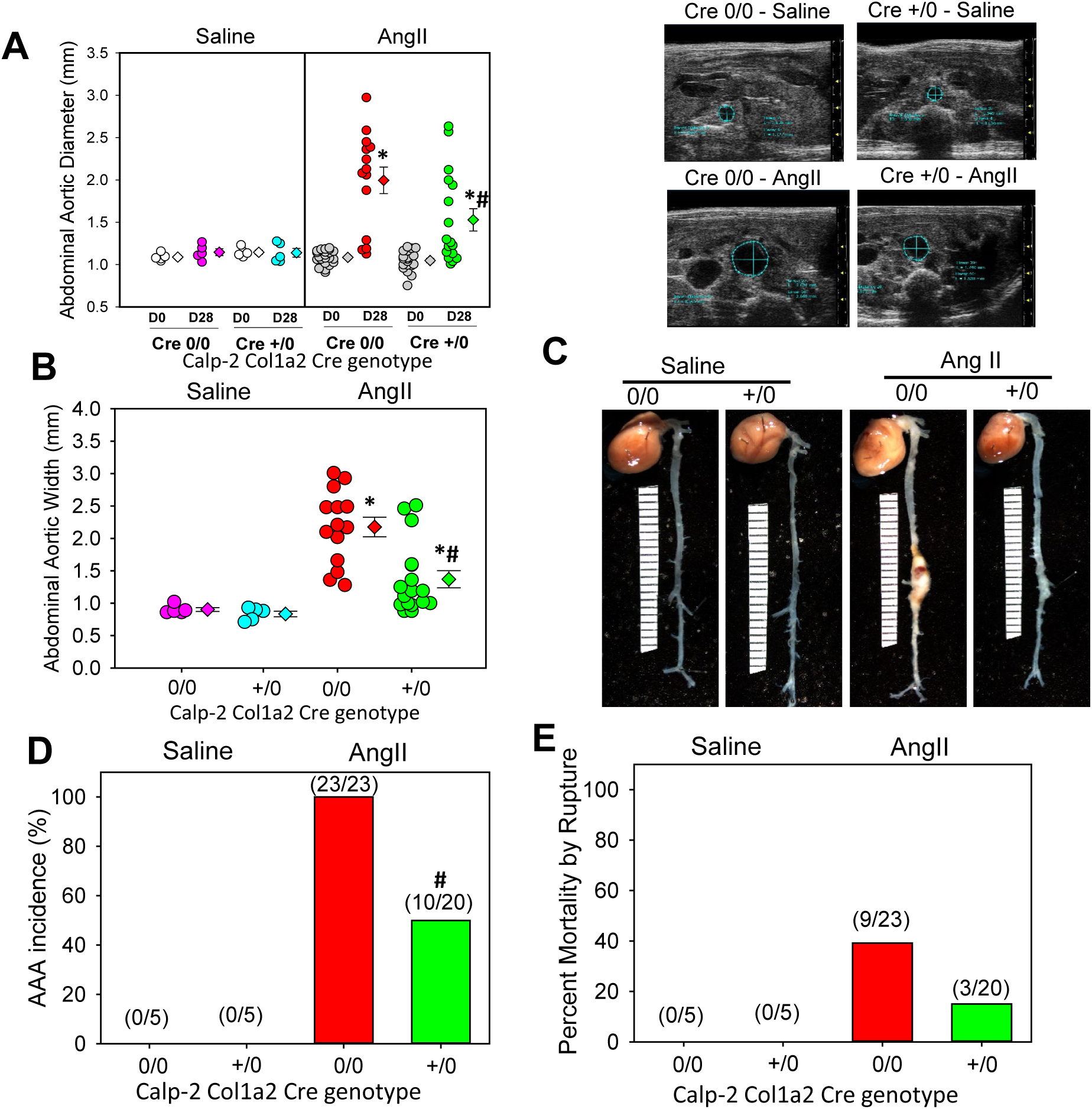
Calpain-2 deficiency in Col1a2 positive mesenchymal cells reduced AngII-induced AAA in Mice. **A**. Ultrasonic measurements of abdominal aortic luminal diameters were measured on day 0 and after 28 days of saline or AngII infusion (Saline n=5; AngII n=20-23). **B,C**. Measurements of maximal external width of abdominal aortas (Saline n=5; AngII n=20-23). Pink (saline - Cre 0/0), teal (saline - Cre +/0), red (AngII - Cre 0/0) and green (AngII - Cre +/0) represent individual mice, diamonds represent means, and bars are SEMs. * denotes P<0.05 when comparing saline vs AngII infusion; # denotes P<0.05 when comparing Cre 0/0 vs Cre +/0 mice (Two-way ANOVA with Holm-Sidak post hoc analysis). **D**. The incidence of AAA (>50% increase in aortic width) in AngII-infused calpain-2 f/f mice that were either Col1a2 Cre 0/0 (red bar) or Cre +/0 (green bar). Statistical analyses were performed by Fisher Exact test (# denotes P<0.001 when comparing Cre 0/0 vs Cre +/0 mice). **E**. Mortality due to AAA rupture in AngII-infused calpain-2 f/f mice that were either Col1a2 Cre 0/0 or +/0. Red (Cre0/0) and green bar (Cre+/0).

### Fragmentation of Cytoskeletal Structural Proteins are Increased in Human AAAs

In AAA tissues, structural integrity of SMC rich medial tissue is highly disrupted (2-4) and many studies have focused extensively on extracellular matrix (ECM) degradation as a part of the mechanism.(25-27) Currently, no reports are available on the cytoskeletal structural proteins which maintain cellular structural integrity by connecting contractile proteins, α-actin and myosin, with the ECM. Filamin A, an actin binding protein, and talin, an integrin binding protein, contribute to organization and stability of the actin cytoskeleton, integrate cellular signaling cascades, and regulate cellular functions including adhesion and motility.(28-30) To examine whether human AAA is associated with fragmented cytoskeletal proteins, tissue lysates from human AAA and control aortas were analyzed for filamin A and talin fragmentation by Western blot using antibodies specific for their C-terminal domain. C-terminal fragmentation of filamin A and talin (Figure 6A,B) were significantly increased in human AAA compared to control aortic tissue, without influencing intact filamin A and talin. These data clearly demonstrate that increased cytoskeletal structural linker protein fragmentation is strongly associated with human AAAs.

**Figure 6.**
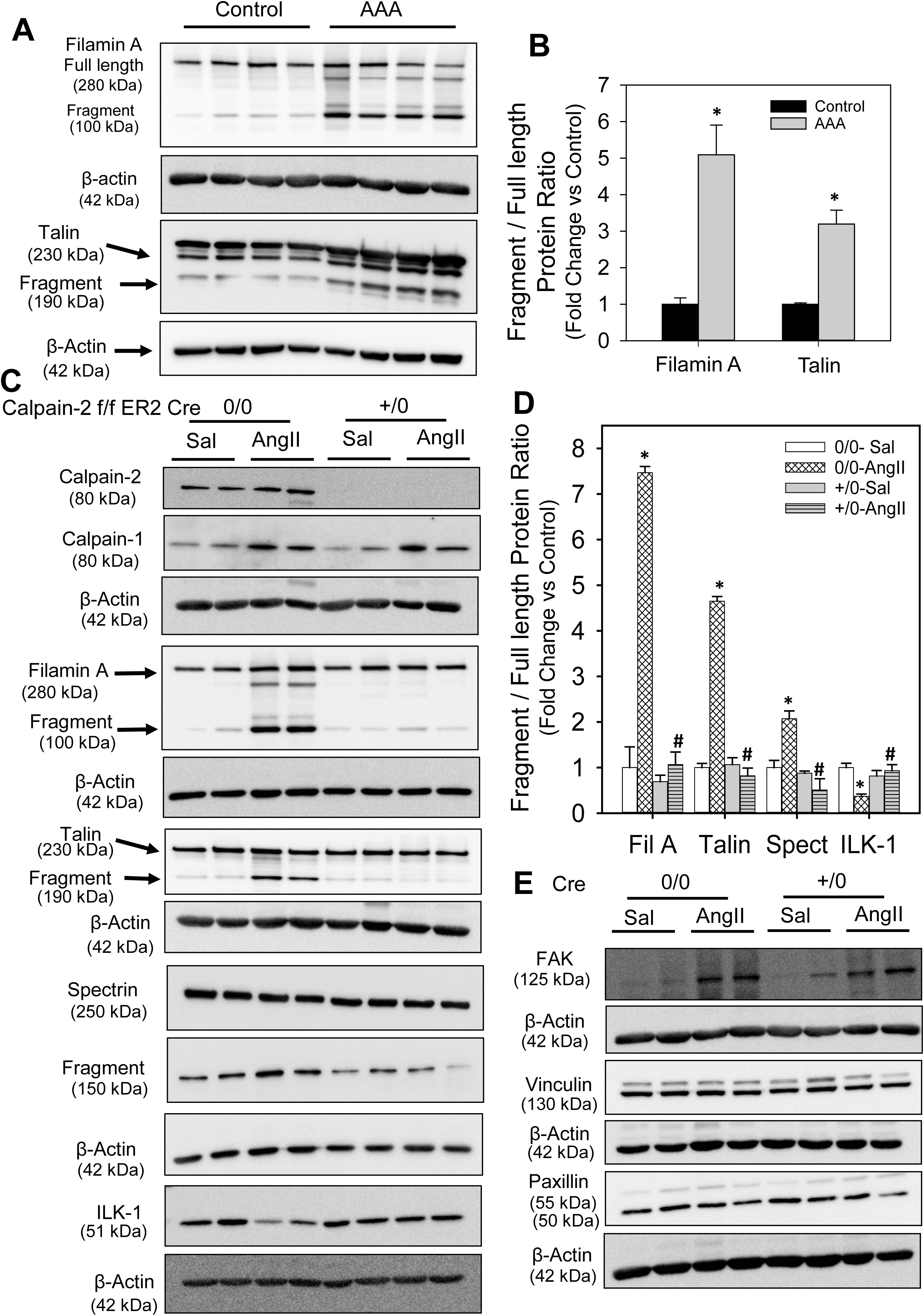
Fragmentation of cytoskeletal structural proteins is increased in human AAA tissues. **A,B**. Full length and fragments of filamin A and talin proteins were detected in human control abdominal aorta and AAA tissues (n=4). * denotes *P*<0.05 when comparing control vs AAA (Student’s *t* test or Mann-Whitney Rank Sum test). **Calpain-2 deficiency significantly suppressed AngII-induced cytoskeletal structural protein fragmentation. C,D**. Full length and fragments of filamin A, talin and spectrin proteins, and ILK-1 and β-actin proteins were detected in abdominal aortas from saline and AngII-infused calpain-2 f/f β-actin Cre0/0 and +/0 mice (n=4). **E**. Focal adhesion kinase (FAK), vinculin, paxillin, and β-actin proteins were detected in abdominal aortas from saline and AngII-infused calpain-2 f/f β-actin Cre0/0 and +/0 mice (n=4). * denotes *P*<0.05 when comparing saline vs AngII infusion; # denotes *P*<0.05 when comparing Cre 0/0 vs Cre +/0 mice (Two-way ANOVA with Holm-Sidak post hoc analysis).

### Calpain-2 deficiency Suppressed AngII-induced Cytoskeletal Structural Protein Fragmentation in the Abdominal Aorta

To examine if calpain-2 regulates cytoskeletal structural protein integrity, first we tested whether AngII infusion promoted cytoskeletal structural protein fragmentation in aortas of mice. Calpain-2 f/f mice that were β-actin Cre0/0 or +/0 were fed a saturated fat-enriched diet and infused with either saline or AngII for 7 days. Western blot analyses using a calpain-2 specific antibody demonstrated that AngII-infusion increased calpain-2 protein abundance in calpain-2 f/f Cre 0/0 mice and none in Cre+/0 aortas as expected, with further phenotypic analysis confirming the deficient genotype (Figure 6C). Similarly, Western blotting using a calpain-1 specific antibody showed that AngII infusion increased calpain-1 protein abundance similarly in both Cre0/0 and +/0 mice with no further compensatory increase in the absence of calpain-2 (Figure 6C). In addition, calpain-2 deficiency significantly suppressed AngII-induced calpain activity in the abdominal aorta (Supplemental Figure S8). Similar in human AAAs, AngII infusion significantly promoted fragmentation of cytoskeletal proteins, filamin A, talin and spectrin, in the abdominal aorta of mice (Figure 6 C,D). Interestingly, calpain-2 deficiency completely prevented the effect of AngII-induced fragmentation on filamin A, talin and spectrin. In addition, calpain-2 deficiency also significantly prevented the AngII-induced suppression of integrin linked kinase (ILK-1; Figure 6C,D), which plays an essential role in connecting the cytoplasmic tail of β subunits of integrins to the actin cytoskeleton, and regulating actin polymerization.(31) However, calpain-2 deficiency had no influence on AngII-induced focal adhesion kinase protein (Figure 6E). In addition, protein abundance of other cytoskeletal proteins such as vinculin and paxillin were not influenced by either AngII or calpain-2 deficiency in the aorta (Figure 6E). Consistent with the aortic tissue, siRNA mediated silencing of calpain-2 in human (Figure 7A) or rat (Figure 7D) aortic SMCs also prevented AngII-induced filamin A (Figure 7B, E) and talin (Figure 7C, F) fragmentation which further confirms the contribution of calpain-2 in promoting fragmentation of cytoskeletal structural proteins, e.g. filamin A and talin.

**Figure 7.**
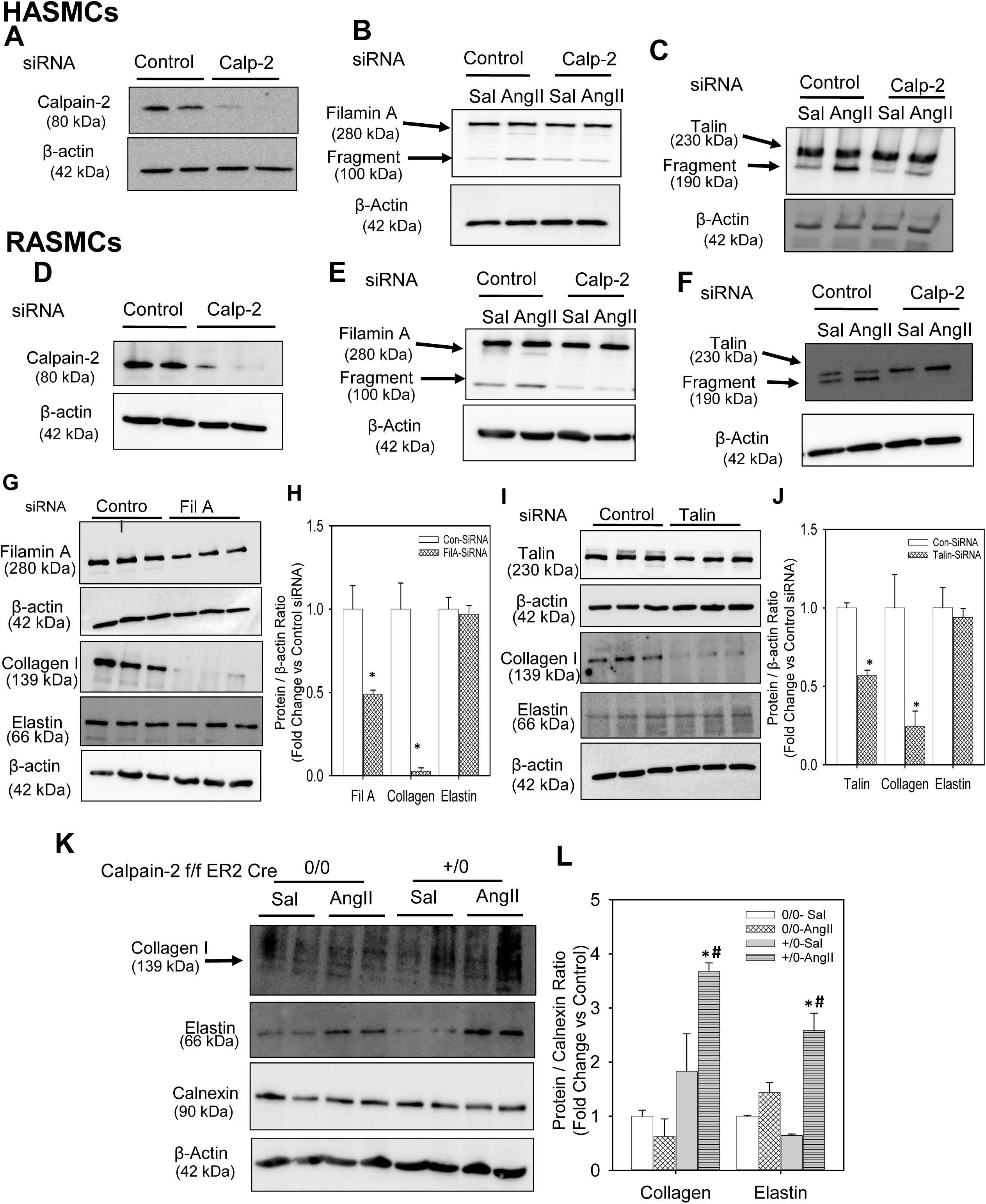
Calpain-2 silencing suppressed AngII-induced filamin A and talin fragmentation in aortic SMCs. Calpain-2 and β-actin proteins were detected in control or calpain-2 siRNA transfected human (**A**) and rat (**D**) aortic SMCs. Full length and fragment of filamin A (**B,E**) and talin (**C,F**) were detected in control or calpain-2 siRNA transfected human and rat aortic SMCs incubated with saline or AngII (1µM or 100 nM for 24 h; n=4). Filamin A, collagen I, elastin and β-actin protein were detected in control, filamin A (**G,H**) or talin (**I,J**) siRNA transfected rat aortic SMCs. * denotes *P*<0.05 when comparing control vs filamin A or talin siRNA. (Student’s *t* test). Collagen I, elastin, calnexin and β-actin protein were detected in abdominal aortas from saline and AngII-infused calpain-2 f/f β-actin Cre0/0 and +/0 mice (n=4; **K,L**). * denotes *P*<0.05 when comparing saline vs AngII infusion; # denotes *P*<0.05 when comparing Cre 0/0 vs Cre +/0 mice (Two-way ANOVA with Holm-Sidak post hoc analysis).

### Silencing of Filamin A or Talin Decreased ECM Protein, Collagen I in Aortic SMCs

Cytoskeletal structural proteins act as linker proteins that hold contractile filaments, actin or myosin, intact with extracellular matrix proteins. (5-8) In order to understand and demonstrate whether loss of cytoskeletal proteins had any influence on ECM proteins, we tested the effect of siRNA mediated silencing of filamin A or talin on ECM proteins, collagen and elastin. Transfection of rat aortic SMCs with either filamin A or talin siRNA for 48 h showed a significant reduction of filamin A (50%) and talin (45%) protein compared to control siRNA transfected cells. Western blot analyses using antibodies specific against collagen I or elastin demonstrated that silencing of either filamin A (Figure 7 G, H) or talin (Figure 7 I, J) strongly suppressed collagen I protein (Figure 7 G,H, I,J) in aortic SMCs; whereas it had no effect on elastin protein (Figure 7 G, H, I, J). Similarly, in mouse aortic SMCs transfected with filamin A or elastin siRNA, immunofluorescent staining using antibodies specific against collagen I revealed a strong reduction of collagen protein (Supplemental Figure S10 and S11).

In addition, to understand the effect of calpain-2 deficiency on aortic collagen and elastin, we performed Western analyses using abdominal aortic tissue lysates from Calpain-2 f/f β-actin Cre0/0 and Cre+/0 mice. AngII infusion for 7 days showed no change on collagen I protein in Cre0/0 groups; whereas it showed a significant increase in collagen I protein in Cre+/0 group (Figure 7 K, L). In addition, interestingly, calpain-2 deficiency showed a significant increase in aortic elastin protein upon AngII infusion compared to Cre0/0 group aortas (Figure 7 K, L).

### Calpain-2 Deficiency Suppressed Rupture of Established AAAs

Several studies have focused on ECM degradation(25) by extracellular proteases such as matrix metalloproteinases (MMPs) (26, 32, 33) and cathepsins (27, 34, 35) as a mechanism; however, presently, no reports prove that inhibition of these proteases are beneficial in limiting AAA progression. Studies have demonstrated that prolonged infusion of AngII to hyperlipidemic mice resulted in progressive dilation of the aortic lumen,(36) which suggest that tamoxifen-inducible deletion of calpain-2 after an AAA formed from AngII infusion may be used to assess the effect of calpain-2 deficiency on AAA progression.

To examine whether calpain-2 deficiency had any influence on the progression of established AAAs, a set of calpain-2 f/f β-actin Cre0/0 and Cre+/0 mice were infused with saline or AngII for 28 days. The ultrasound measurement of aortic luminal dilation was performed on day 0 and 21 to ensure the development of AAAs (Figure 8 A, B, C). On day 23, the mice were injected with tamoxifen for 5 consecutive days to activate the Cre recombinase (Figure 8 A). The mice were continuously infused with saline or AngII for another 8 weeks and the aortic luminal dilation were periodically measured at days 42 and 70 of infusion (Figure 8 D, F). On day 84, the mice were terminated and the aortas were examined. Inducible depletion of calpain-2 did not influence the progressive lumen dilation (Figure 8 F, G) and maximal ex-vivo aortic width expansion of AngII-induced established AAAs (Figure 8 H, I) and AAA incidence (Figure 8 J). Interestingly, inducible depletion of calpain-2 significantly suppressed the mortality induced by aortic rupture (Figure 8 K) and improved the survival of calpain-2 deficient mice compared to control from AngII-induced aortic rupture mediated death (Figure 8 L).

**Figure 8.**
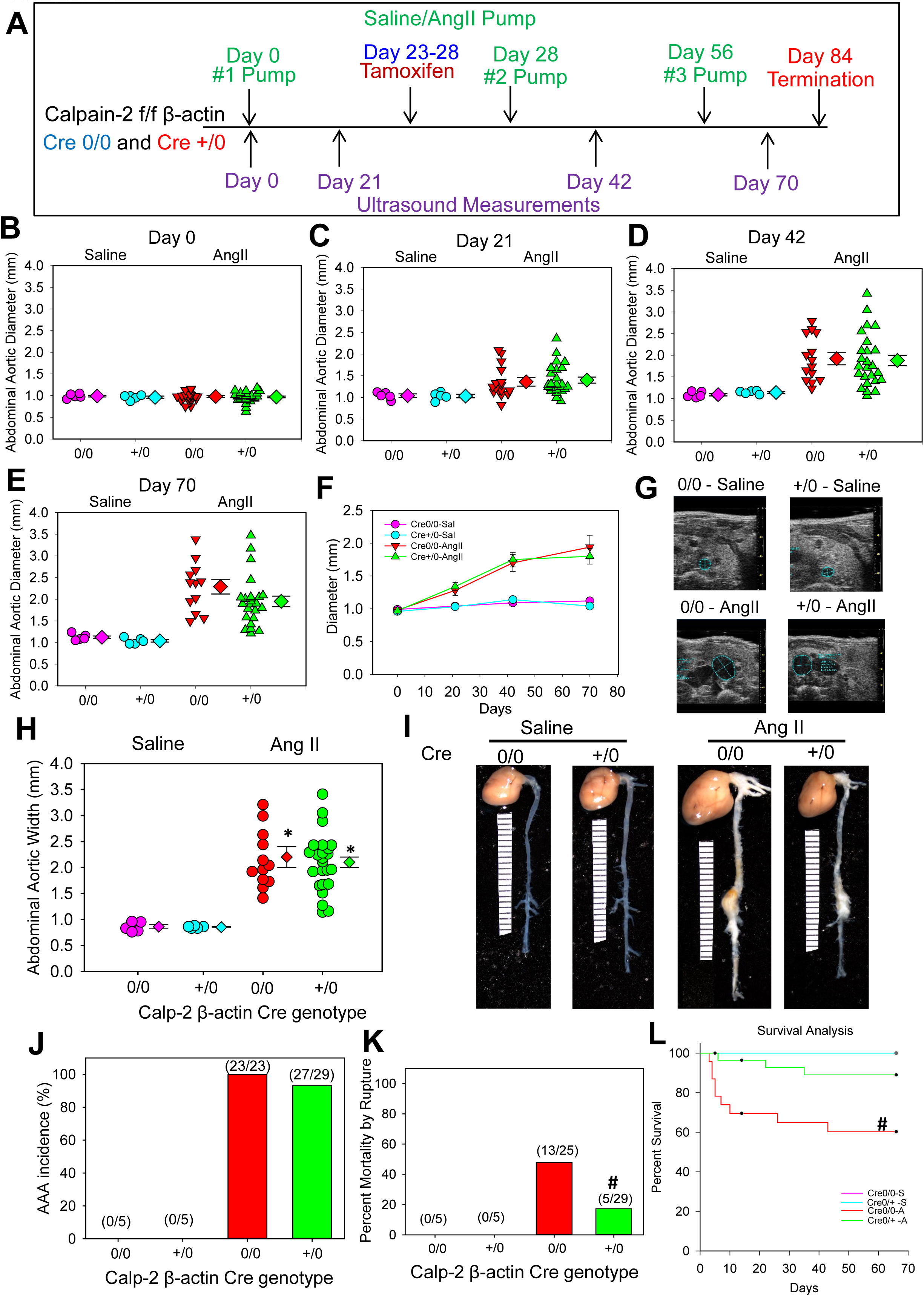
Inducible depletion of Calpain-2 deficiency significantly suppressed rupture of established AAAs. **A**. Experimental design to study the effect of inducible-calpain-2 deficiency on the progression of AngII-induced established AAAs. Ultrasonic measurements of abdominal aortic luminal diameters were measured on day 0 **(B)**, after 21**(C)**, 42 **(D)**, and 70 **(E)**, days of saline or AngII infusion (Saline n=5; AngII n=23-29). **F**. Representation of progressive luminal dilation over the course of 84 days of saline or AngII infusion. **G**. Representative images of ultrasound pictures of abdominal aortas of Cre 0/0 and +/0 mice infused with saline or AngII. **H, I**. Measurements of maximal external width of abdominal aortas (Saline n=5; AngII n=23-26). Pink (saline - Cre 0/0), teal (saline - Cre +/0), red (AngII - Cre 0/0) and green (AngII - Cre +/0) represent individual mice, diamonds represent means, and bars are SEMs. * denotes P<0.05 when comparing saline vs AngII infusion; # denotes P<0.05 when comparing Cre 0/0 vs Cre +/0 mice (Two-way ANOVA with Holm-Sidak post hoc analysis). **J**. The incidence of AAA (>50% increase in aortic width) in AngII-infused calpain-2 f/f mice that were either β-actin Cre 0/0 (red bar) or Cre +/0 (green bar). **K**. Mortality due to AAA rupture in AngII-infused calpain-2 f/f mice that were either β-actin Cre 0/0 or +/0. Red (Cre0/0) and green bar (Cre+/0). Statistical analyses were performed by Fisher Exact test (# denotes P<0.001 when comparing Cre 0/0 vs Cre +/0 mice). **L**. Survival rate from the abdominal aortic rupture.

## DISCUSSION

In the present study, first we showed the clinical relevance of calpain-2 to AAA by demonstrating the presence of increased calpain-2 protein and activity in human AAA tissues. In addition, we demonstrated that human AAAs are associated with increased cytoskeletal structural protein fragmentation. Furthermore, to determine the contribution of calpain-2 in AAA formation and cytoskeletal structural proteins in the aorta, we generated LDL receptor -/- mice with tamoxifen-inducible whole body calpain-2 deficiency using calpain-2 floxed and transgenic mice that express Cre recombinase under the control of the ubiquitous promoter, chicken β-actin. Using this unique calpain-2 deficient mouse model, we examined the role of calpain-2 in AngII-induced AAAs. Here, we demonstrated that inducible depletion of calpain-2 in adult mice significantly reduced AngII-induced AAA formation. The beneficial effect of calpain-2 deficiency on AngII-induced AAA was associated with strong reduction of cytoskeletal structural protein destruction in the abdominal aorta of mice. Furthermore, using tamoxifen-inducible FMSC (including SMCs and fibroblasts) and adipocyte-specific calpain-2 deficient mice, we demonstrated that calpain-2 derived from vessel wall cells not from surrounding perivascular adipose tissue (PVAT) contributes to AngII-induced AAA development. In addition, in cultured aortic SMCs, we demonstrated that silencing of cytoskeletal structural proteins, filamin A or talin, significantly reduced abundance of collagen protein which supports a functional role of cytoskeletal structural proteins for ECM protein integrity.

Calpain-2 deficiency did not have any effects on AngII-induced blood pressure elevation. Consistent with our current observation, our earlier studies of calpain inhibition by BDA-410, or calpain-2 deficiency in leukocytes also showed no effect on AngII-induced increased blood pressure.(15, 17) In addition, development of AngII-induced AAA is independent of increases in blood pressure.(37)

Whole body inducible depletion of calpain-2 significantly reduced AngII-induced AAA. Our study clearly demonstrates that depletion of calpain-2 is sufficient enough to significantly suppress AngII-induced calpain activity in the abdominal aorta. In our earlier studies, we showed that deficiency of calpain-1, another major isoform of calpain, or calpain-2 deficiency in leukocytes had no effect on AngII-induced AAA.(16, 17) Our current observation clearly demonstrates that calpain-2 derived from vessel wall cells (SMCs/ fibroblasts) plays a critical role in AngII-induced AAA formation in mice. Fibrogenic mesenchymal cells that including fibroblasts, specialized myofibroblasts, synthetic aortic SMCs, which in common share the phenotype of expressing Col1, play a critical role in AngII-induced aortic fibrosis and ECM remodeling.(10) A series of studies highlighted that these fibrogenic mesenchymal cells regulates ECM remodeling through MMP activation,(38) fibronectin production,(39) vascular inflammation via RelA/NF-kB signaling and cytokine production.(10) To our knowledge, this is the first study to report the functional contribution of intracellular protease, calpain-2 in the regulation of ECM in fibrogenic mesenchymal cells, during AAA development. In support, in our study, immunohistochemical staining showed a strong distribution of calpain-2 in fibroblast rich adventitia and a week distribution in SMC rich media in addition to infiltrating leukocytes and surrounding PVAT of human and AngII-induced AAA. Our earlier study using leukocyte-specific calpain-2 deficient mice (17) and our current observation using adipocyte-specific clapin-2 deficient mice, clearly demonstrates that calpain-2 derived from fibrogenic mesenchymal cells (SMCs/ fibroblasts), but not from infiltrating leukocytes or surrounding PVAT plays a critical role in AngII-induced AAA formation in mice.

Our current study is the first to report the presence of cytoskeletal structural protein fragmentation in both human and experimental AAAs. Cytoskeletal structural proteins act as linker proteins that hold contractile filaments, actin or myosin, intact with extracellular matrix proteins. (5-8) Loss of these structural protein may lead to destruction of cell structural integrity. Calpain-2 deficiency or silencing completely prevented AngII-induced fragmentation of filamin A and talin in mice and cultured SMCs. In addition, gene silencing of filamin A or talin in cultured aortic SMCs lead to complete loss of ECM protein collagen, but had no effect on elastin. However, the differential regulation of collagen and elastin by cytoskeletal proteins in cultured aortic SMCs needs further investigation. In addition, our studies using inducible whole body calpain-2 deficient mice clearly showed a significant increase in aortic collagen and elastin during AngII infusion, and significantly suppressed rupture of established AAAs. The data from in vitro and in vivo studies suggest that calpain-2 mediates AngII-induced cytoskeletal structural protein destruction, which in turn may promote ECM disorganization and disruption of aortic structural integrity. These observations clearly indicate that cytoskeletal structural proteins play a critical role in maintaining ECM proteins. Further studies are warranted to understand the mechanisms by which cytoskeletal structural proteins help to maintain the stability of ECM.

Currently, it is not clear by which mechanism calpain-2 mediates or contributes to cytoskeletal structural protein destruction. Calpain cleavage site prediction analyses using the online calpain cleavage site predictor software (40) (CaMPDB calpain.org) revealed various strong cleavage sites in structure of filamin A and talin. Filamin A has a calpain binding site at Ser2152 residue in the H1 region.(41) Similarly talin contains two calpain-2 binding sites at Leu432 and Glu2492.(13, 42, 43) However, further studies are warranted to understand whether calpain-2 mediates its effect on these structural proteins through direct binding on these predicted sites or via an indirect mechanisms. Utilizing cells that express structural protein mutations at potential calpain-2 binding sites, our future studies will delineate the functional role of calpain-2 in cytoskeletal structural protein destruction.

## STUDY LIMITATIONS

Our current study used the genetic approach to evaluate the effect of calpain-2 inhibition on the progression of established AAAs. Further studies are warranted using a calpain-2 specific pharmacological inhibitor and also in complementary experimental animal models from the perspective of using this as a therapeutic approach clinically to halt aortic rupture in humans. However, the results of the present study strongly suggest that inhibition of calpain-2 activity prevent rupture of established AAAs.

## CONCLUSIONS

In summary, we demonstrated that inducible depletion of calpain-2 decreased AAA formation, and rupture of established AAAs, which was associated with reduced cytoskeletal structural protein fragmentation and increased collagen matrix protein in mice (Figure 9). These results suggest that inhibition of calpain-2 may offer a new therapeutic target to reduce AAAs.

**Figure 9.**
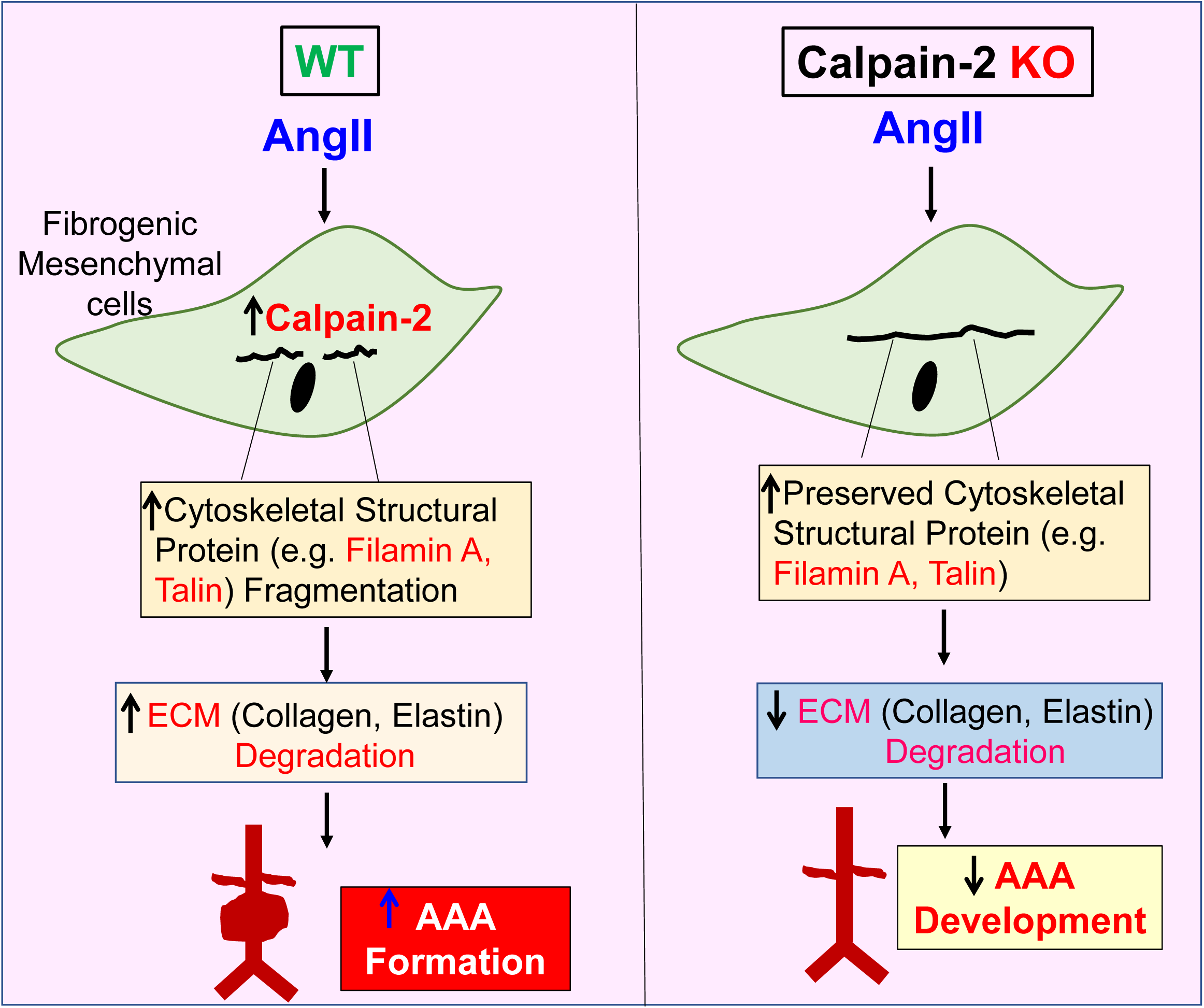
Graphical Summary.

## MATERIALS AND METHODS

### Mice

LDL receptor -/- (stock # 002207), inducible β-actin Cre+/0 (CAG-Cre-ER™; stock # 004682), Col1α2 Cre+/0 (Col1a2-Cre-ER, Stock # 029235), ROSA26R^LacZ^ (Stock # 003474) and C57BL/6J (Stock# 000664) mice were purchased from The Jackson Laboratory (Bar Harbor, ME). Calpain-2 floxed (f/f) mice on a C57BL/6 background were originally generated in the laboratory of Dr. Takaomi Saido.(18) LDL receptor -/- and calpain-2 f/f mice were backcrossed 10 times into a C57BL/6 background.

Both calpain-2 f/f and β-actin/ Col1α2 Cre+/0 mice were bred to an LDL receptor -/- background. Female calpain-2 f/f mice were bred with male Cre+/0 mice to yield mice homologous for the floxed calpain-2 gene and hemizygous for the Cre transgene. Littermates that were homozygous for the floxed calpain-2 gene, but without the Cre transgene (Cre0/0), were used as control mice.

Age-matched male littermates (8-10 weeks old) were used for the present study. Mice were maintained in a barrier facility and fed normal mouse laboratory diet. All study procedures were approved by the University of Kentucky Institutional Animal Care and Use Committee (Protocol # 2011-0907). This study followed the recommendations of The Guide for the Care and Use of Laboratory Animals (National Institutes of Health).

### Mouse Genotyping

Mouse genotypes were confirmed by PCR. DNA was isolated from tail snips using a Maxwell tissue DNA kit (Cat# AS1030, Promega, Madison, WI). Calpain-2f/f genotyping used the following primers: 5’-ATAGCTCCTGTGTATCAG GCACAGAGCTGG-3’ and 5’-CTCTGGTCAGGTCTTAGTTCCCAGAGGATG - 3’. Resultant wild-type, and flox allele bands were 290, and 430 bp, respectively (Supplemental Figure S1A). Cre+ genotyping used the following primers: 5’-ACCTGAAGATGTTCGCGATT and 5’-CGGCATCAACGTTTTCTTTT. The resultant Cre+ hemizygous allele PCR product was 182 bp and no product for non-transgenic mice. The IL-2 gene was used as an internal control for Cre+ genotyping using the following primers: 5’-CTAGGCCACAGAATTGAAAGAT CT and 5’-GTAGGTGGAAATTCTAGCATCA TCC. The resultant product was 324 bp (Supplemental Figure S1B). LDL receptor genotyping was performed as described previously.(44)

### Tamoxifen Injections

Tamoxifen (Cat# T5648; Sigma Chemical; St. Louis, MO) was dissolved in 100% ethanol (30 mg in 300 µl) and then diluted in corn oil (Cat# C8267; Sigma Chemical; St. Louis, MO) for a final concentration of 10 mg/ml. Mice (6-8 weeks old) were injected i.p. with a dose of tamoxifen of 25 mg/kg daily for 5 consecutive days.

### Diet

To induce hypercholesterolemia, mice were fed a diet supplemented with saturated fat (21% wt/wt milk fat; TD.88137, Harlan Teklad, Indianapolis, IN) for 5 weeks.

### AngII Infusion

Two weeks after last tamoxifen injection and an initial week of high-fat diet feeding, mice were implanted with Alzet osmotic minipumps (Model 2004, Durect Corporation, Cupertino, CA), subcutaneously into their right flanks, and infused with either saline or AngII (1,000 ng/kg/min, Bachem, Torrance, CA) continuously for a period of 7 or 28 days, as described previously.(16, 45) Mice were fed a high fat-enriched diet throughout the infusion study.

### Blood Pressure Measurement

Systolic blood pressure (SBP) was measured noninvasively on conscious mice by volume pressure recording of the tail using a computerized tail cuff blood pressure system (Kent Scientific Corp, Torrington, CT).(46) SBP was measured on 5 consecutive days prior to pump implantation, and during the last 5 days of AngII infusion.

### Measurement of Plasma Components

Plasma cholesterol concentrations were measured using a commercially available enzymatic kit (Wako Chemicals, Richmond, VA) as described previously.(44) Plasma renin concentrations were measured by generation of Angiotensin I during incubation of plasma as described previously. (47)

### Ultrasound Imaging of Abdominal Aortic Aneurysms

Luminal dilation of the abdominal aorta was measured by a high frequency ultrasound imaging system (Vevo 2100, Visual Sonics, Toronto, Canada) using a MS400 MicroScanTM transducer with a resolution frequency of 18–38 MHz.(48) Mice were anesthetized and restrained in a supine position to acquire ultrasonic images. Short axis scans of abdominal aortas were performed from the left renal arterial branch level to the suprarenal region.(48) Images of abdominal aortas were acquired and measured to determine maximal diameter in the suprarenal region of the abdominal aorta. Aortic images were acquired at day 0 and 28 of AngII-infusion.

### Quantification of Atherosclerosis and Abdominal Aortic Aneurysms

After saline was perfused through the left ventricle of the heart, aortas were removed from the origin to iliac bifurcation, and placed in formalin (10% wt/vol) overnight. Adventitial fat was cleaned from the aortas. Atherosclerosis was quantified on aortic arches and thoracic aorta as lesion area, and percent lesion area on the intimal surface by en face analysis as described previously.(49-51) Lesion areas were measured using Image-Pro Plus software (Media Cybernetics, Bethesda, MD) by direct visualization of lesions under a dissecting microscope. For aneurysm measurements, AAAs were quantified *ex vivo* by measuring the maximum external width of the suprarenal abdominal aortic diameter using computerized morphometry (Image-Pro Cybernetics, Bethesda, MD) as described previously.(16, 52)

### Tissue Histology

Abdominal aortas were placed in optimal cutting temperature (OCT) compound and serially sectioned (10 μm thickness/section) in sets of 10 slides with 9 sections/slide using a cryostat.(53) One slide of each serial set was stained with Movat’s Pentachrome (Polyscientific) to visualize elastin breaks.

### β-Galactosidase staining in tissues

Whole tissues were fixed in paraformaldehyde for 1 hour, and preincubated with buffer containing100 mM sodium phosphate, pH7.3; 2 mM magnesium chloride; 0.01% sodium deoxycholate; 0.02% NP-40 (Cat# S-3139; M-8266; D6750; 74385; Sigma-Aldrich; St. Louis, MO, respectively). Subsequently, tissues were incubated overnight in staining buffer (above buffer with additions of 5 mM potassium ferricyanide, 5 mM potassium ferrocyanide; Cat # 702857; P3289; Sigma-Aldrich; St. Louis, MO, respectively) containing β-galactosidase (1 mg/ml; Cat # V394A; Promega, Madison, WI). Samples were post fixed in formalin overnight and then stored in ethanol (70%).

### Immunohistochemistry of Human AAAs

Calpain-2 immunostaining was performed using a rabbit anti-human calpain-2 (10 µg/ml, catalog No. RP-3 Calpain-2; Triple Point biologics, Forest Grove, OR). CD68 immunostaining was performed using a rabbit anti-human CD68 antibody (1:200, catalog No. SC-9139; Santa Cruz Biotechnology, Dallas, TX). Biotinylated secondary antibodies from the appropriate species were used (Vector Laboratories). A peroxidase-based ABC system (Vector Laboratories) and the red chromogen AEC were used to detect the antigen-antibody reaction. Immunostaining was performed on formalin-fixed paraffin-embedded sections, with appropriate negative controls, as described previously.(44, 54) Human control abdominal and AAA sections were surgical samples procured by Dr. John Curci at Washington University, MO. The sample details were:

| <u>Sample</u> | <u>Age</u> | <u>Gender</u> |
| --- | --- | --- |
| Control abdominal aorta -1 | 56 | M |
| Control abdominal aorta -2 | 64 | F |
| Control abdominal aorta -3 | 51 | M |
| AAA-1 | 61 | M |
| AAA-2 | 70 | F |
| AAA-3 | 68 | M |

### Western Blot Analyses

#### Mouse Aortic Lysate

Abdominal aortic tissue lysates were extracted in radioimmunoprecipitation assay lysis buffer, and protein content was measured using a Bradford assay (Bio-Rad, Hercules, CA). Protein extracts (20-30 μg) were resolved by SDS-PAGE (6.0 or 7.5 % wt/vol) and transferred electrophoretically to PVDF membranes (Millipore). After blocking with non-dry fat milk (5 % wt/vol), membranes were probed with primary antibodies. The following antibodies were used (please see the Major Resources Table in the Supplemental Material): calpain-1 domain IV (Abcam, catalog No: ab39170), calpain-2 (Abcam, catalog No: ab39165), filamin A (Abcam, catalog No: ab76289), talin (Abcam, catalog No: ab71333), α-spectrin (Millipore, catalog No: MAB1622), ILK-1 (Cell Signaling, catalog No:3862), FAK (Cell Signaling, catalog No:3009), vinculin (Abcam, catalog No: ab73412), paxillin (Abcam, catalog No: ab32264), collagen (Abcam, catalog No: ab34710), elastin (Abcam, catalog No: ab217356), calnexin (Enzo, catalog No: ADI-SPA-860-F) and β-actin (Sigma-Aldrich, catalog No: A5441). Membranes were incubated with appropriate HRP-labeled secondary antibodies. Immune complexes were visualized by chemiluminescence (Pierce, Rockford, IL) and quantified using a Kodak Imager.

#### Human AAA Lysate

Human AAA tissue were fresh frozen surgical samples procured by Dr. Gerard Pasterkamp, at the University Medical Center Utrecht, Netherlands. As controls, fresh frozen human infra-renal aortic tissues were purchased from the Tissue for Research Company, UK. These control tissues were collected within 3 h post-mortem from organ donors in a deidentified manner. The sample details are:

| <u>Sample</u> | <u>Age</u> | <u>Gender</u> |
| --- | --- | --- |
| Control abdominal aorta -1 | 80 | M |
| Control abdominal aorta -2 | 61 | F |
| Control abdominal aorta -3 | 58 | M |
| Control abdominal aorta -4 | 64 | M |
| Control abdominal aorta -5 | 75 | F |
| Control abdominal aorta -6 | 63 | M |
| AAA-1 | 71 | M |
| AAA-2 | 64 | M |
| AAA-3 | 73 | M |
| AAA-4 | 76 | M |
| AAA-5 | 59 | M |
| AAA-6 | 68 | F |

Control or AAA tissue lysates were extracted and protein content was measured in a similar method to mouse aortic tissues. The calpain-2 and β-actin antibodies were used similarly as mentioned above.

### Calpain Activity Assay

Calpain enzyme activity was measured in human AAA and mouse aortic tissue lysates fluorimetrically using a commercially-available activity assay kit (Calpain, catalog No: K240-100; BioVision, Mountain View, CA) as described previously.(15) Aortic protein extracts (20 mg) were incubated with fluorogenic (4-trifluoromethyl coumarin labeled) calpain substrate for 60 min at 37°C. Mean fluorescence signals were measured using a microplate fluorescent plate reader (Spectramax M2; MolecularDevices, Sunnyvale, CA) per manufacturer’s instructions.

### Human Aortic SMCs and Short Interfering RNA Knockdown

Human aortic SMCs were purchased from Promo Cell (Catalog No: C-12533) and cultured in SMC growth medium (Promo Cell, Catalog No: C39262). The SMC cell purity was verified using a SMC α-actin antibody (1:100, Abcam catalog No: ab5694) by immunohistochemistry (Supplemental Figure S9). Cells of passages 2-3 were used for the short interfering (si)RNA transfection study. SMCs were transfected with control siRNA or siRNA targeting the human calpain-2 (Ambion - Silencer Select validated siRNA, Thermo Fisher Scientific) sequences using RNAiMax lipofectamine transfection reagent (Life Technologies, Thermo Fisher Scientific). After 48 h of transfection, cells were incubated with vehicle or AngII (1 µM) for 24 h; then protein lysates were harvested for western blot analyses.

### Rat Aortic SMCs and Short Interfering RNA Knockdown

Rat aortic SMCs were isolated and cultured as described previously.(55) Briefly, rat aortas were excised and dissected free of adventitia and fat. The vessels were further digested by collagenase type I (Worthington Biochemical Corp, 1 mg/ml) and elastase type III (sigma 0.125 mg/ml) at 37°C in a CO_2_ humidified chamber. Then, cell suspensions were centrifuged and the cell pellets were re suspended and grown in DMEM with 10% fetal bovine serum and penicillin and streptomycin (1%) in 5% CO_2_ at 37°C. The SMC phenotype was determined as described above. Cells of passages 2-3 were used for the short interfering (si)RNA transfection study. SMCs were transfected with control siRNA or siRNA targeting the rat calpain-2, filamin A or talin (Ambion - Silencer Select validated siRNA, Thermo Fisher Scientific) sequences using RNAiMax lipofectamine transfection reagent (Life Technologies, Thermo Fisher Scientific). After 48 h of transfection, cells were treated with vehicle or AngII (100 nM) for 24 h for western blot analyses.

### Mouse Aortic SMCs, Short Interfering RNA Knockdown and Immunofluorescence

Mouse aortic SMCs were purchased from Cell Biologics (Catalog No: C57-6080) and cultured in SMC growth medium (Cell biologics, Catalog No: M2268) as described previously.(55) The SMC cell purity was determined as described above. Cells of passages 2-3 were used for the short interfering (si)RNA transfection study. SMCs were transfected with control siRNA or siRNA targeting mouse filamin A, talin or calpain-2 (Ambion - Silencer Select validated siRNA, Thermo Fisher Scientific) sequences using RNAiMax lipofectamine transfection reagent (Life Technologies, Thermo Fisher Scientific). After 48 h of transfection, cells were fixed, permeabilized, and collagen I immunostaining was performed using a rabbit anti-mouse collagen I (1 µg/ml, catalog No. ab34710, Abcam). Fluorescent labelled goat anti-rabbit secondary antibody (1 µg/ml, catalog No. ab1754741, Abcam) was used to visualize positive immunostaining under fluorescent microscopy. The mean fluorescence intensity of 100 cells was measured by Image J software. Fluorescent Cy3-labelled clone 1A4 α-actin antibodies (Clone 1A4, Sigma-Aldrich, catalog No: C6198) were used to visualize cytoskeletal actin protein as described previously.(55) (Please see the Major Resources Table in the Supplemental Material).

### Statistical Analyses

Data are represented as mean ± SEM. Statistical analyses were performed using SigmaPlot 12.0 (SYSTAT Software Inc., San Jose, CA, USA). Repeated measurement data were analyzed with SAS fitting a linear mixed model expressing the temporal trend in systolic blood pressure as a quadratic polynomial in time for each treatment. Student’s *t* test or Mann-Whitney Rank Sum test was performed as appropriate for two-group comparisons. One or Two way ANOVA with Holm-Sidak post hoc analyses were performed for multiple-group and multiple-manipulation analysis. Fisher’s exact probability test was used to determine differences between groups in the incidence of aneurysm formation and in mortality due to rupture. Values of *P*<0.05 were considered statistically significant.

## Abbreviations and Acronyms

AngII: angiotensin II
AAA: abdominal aortic aneurysm
LDL: low-density lipoprotein
ECM: extracellular matrix
SMC: smooth muscle cell

## ACKNOWLEDGMENTS

This study was supported by a Scientist Development Grant (14SDG18740000) from the American Heart Association and by the National Institutes of Health (Grants P20GM103527, R01HL130086). The content in this manuscript is solely the responsibility of the authors and does not necessarily represent the official views of the National Institutes of Health.

## DISCLOSURES

The authors report no conflicts.

## HIGHLIGHTS

- Human and experimental AAAs are associated with increased calpain-2 protein and activity, and increased cytoskeletal structural protein fragmentation.
- Inducible depletion of calpain-2 in mice decreases AAA formation, which is associated with reduced fragmentation of cytoskeletal structural proteins.
- Targeted inhibition of calpain-2 may offer a new therapeutic direction to reduce AAAs.

---

**Figure S1.**
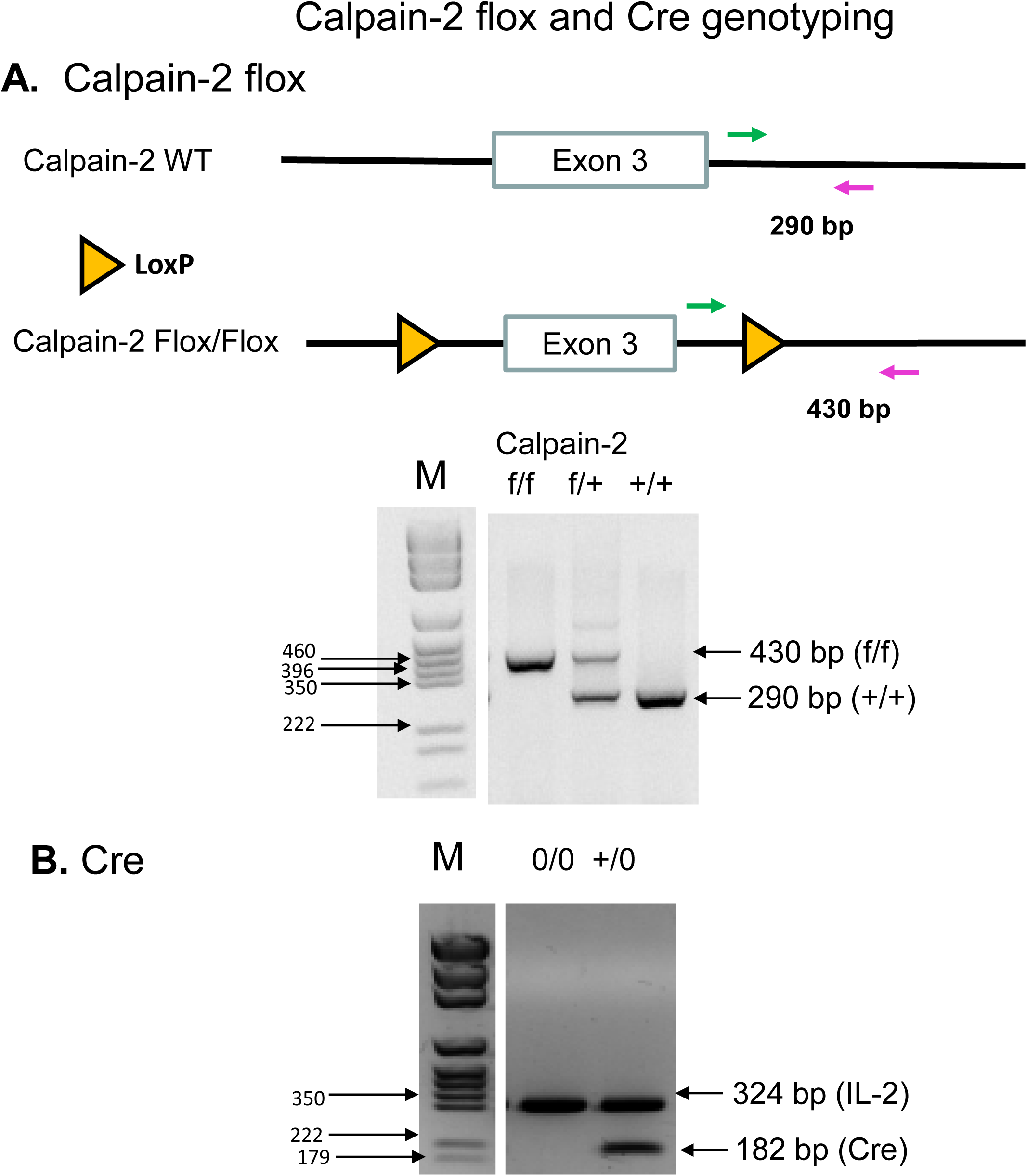
Genotyping of experimental mice for the calpain-2 floxed allele and Cre Tg by PCR. Genomic DNA from tail biopsies was isolated and screened by PCR. **A**. PCR screening strategy to verify the presence of calpain-2 flox/flox. A 430 bp product was generated from calpain-2 f/f mice, while a 290 bp product was obtained from calpain-2 WT mice. **B**. Cre Tg was confirmed by PCR using IL-2 gene as internal control. PCR on genomic DNA yielded amplicons of 182 and 324 bp for Cre Tg and IL-2 alleles, respectively. (M= Molecular weight ladder)

**Figure S2.**
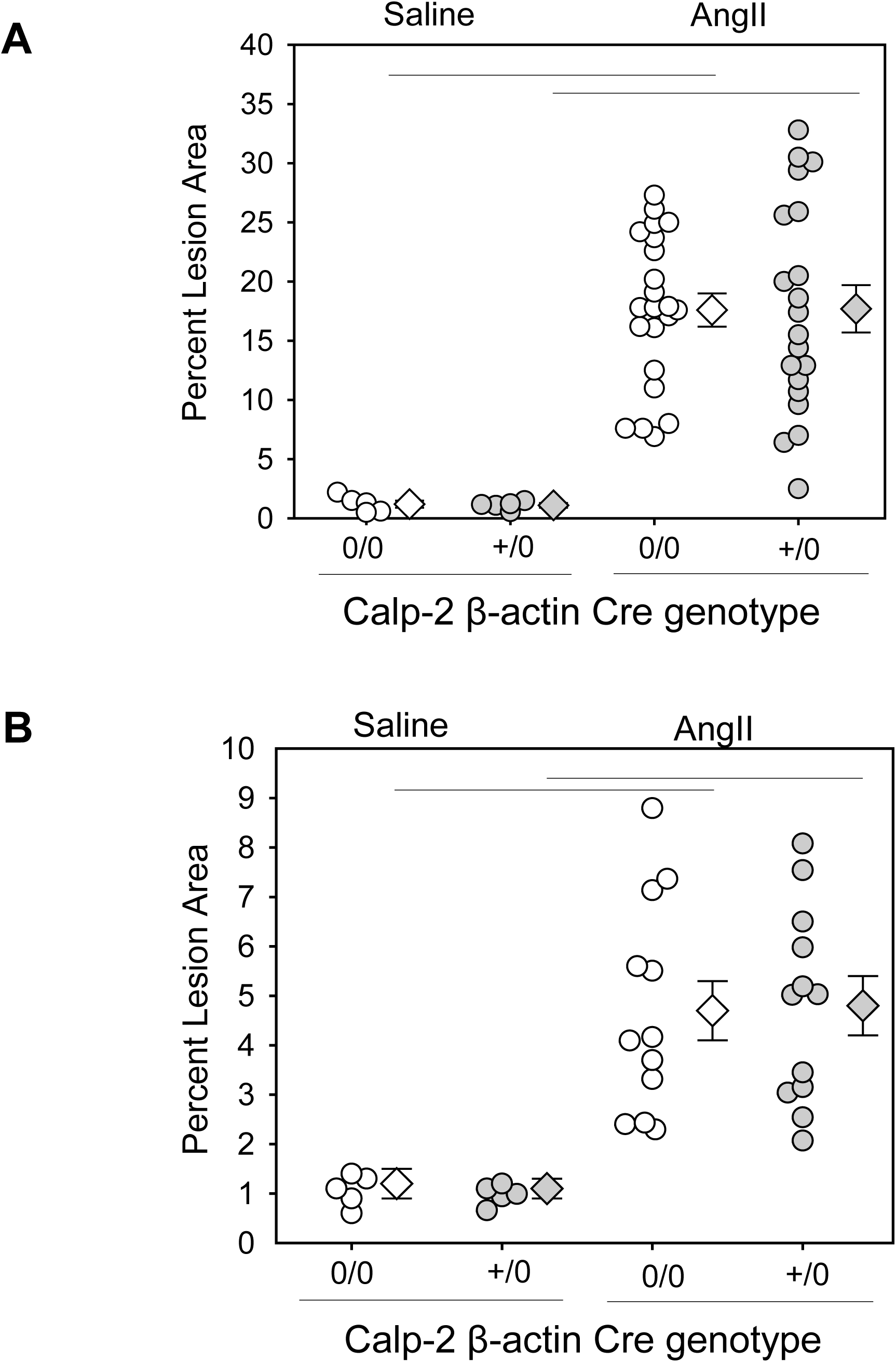
Calpain-2 deficiency had no effect on AngII-induced arch and thoracic atherosclerosis. Atherosclerotic lesion area was measured on the intimal surface of aortic arch (**A**, Saline n=5; AngII n=17-20) and thoracic aorta (**B**, Saline n=5; AngII n=12). White (Cre 0/0) and gray circles (Cre+/0) represent individual mice, diamonds represent means, and bars are SEMs. Horizontal lines represent significance of *P*<0.05 (Two-way ANOVA with Holm-Sidak post hoc analysis).

**Figure S3.**
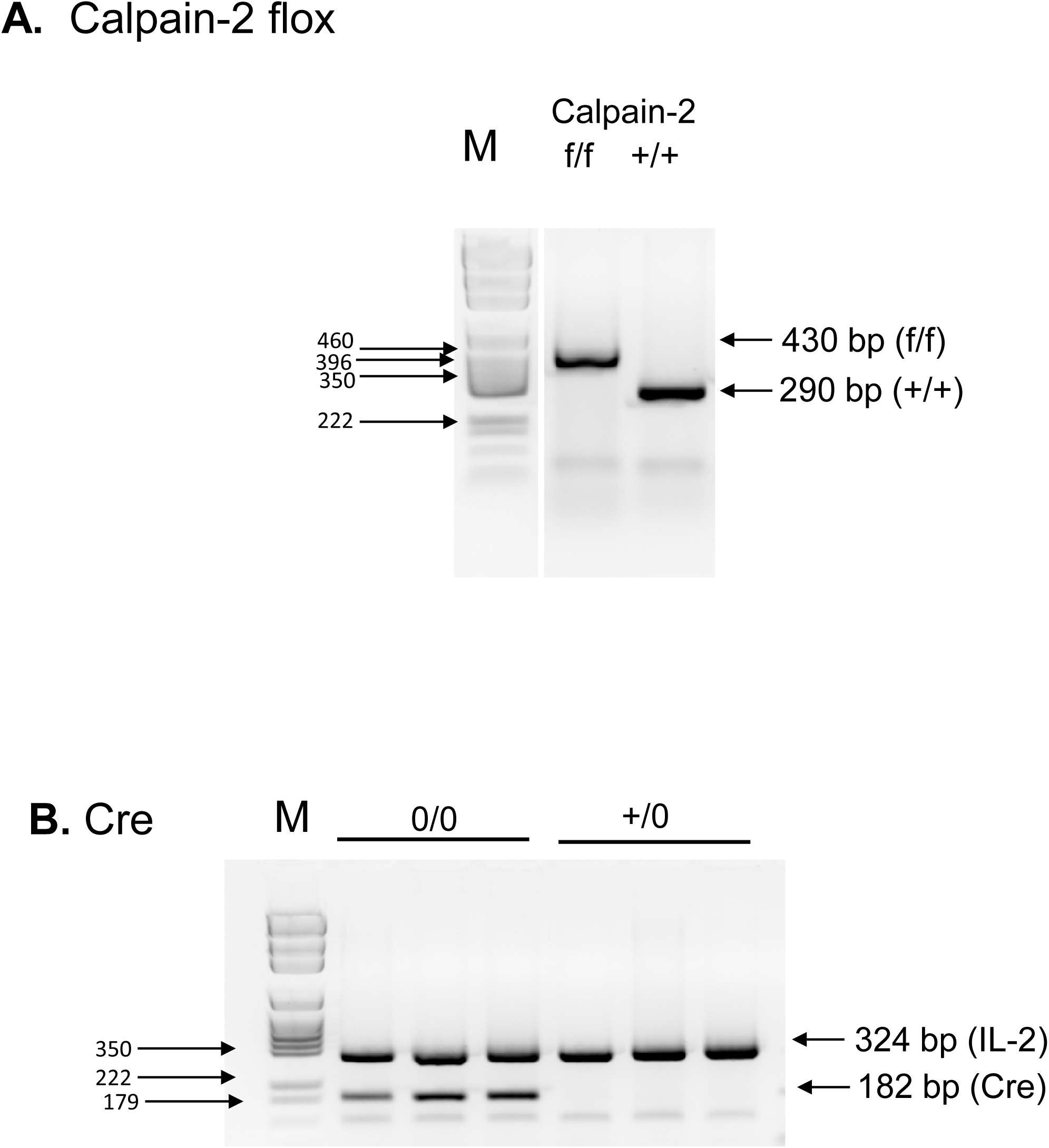
Genotyping of experimental mice for the calpain-2 floxed allele and adiponectin Cre Tg by PCR. Genomic DNA from tail biopsies was isolated and screened by PCR. **A**. A 430 bp product was generated from calpain-2 f/f mice, while a 290 bp product was obtained from calpain-2 WT mice. **B**. Cre Tg was confirmed by PCR using IL-2 gene as internal control. PCR on genomic DNA yielded amplicons of 182 and 324 bp for Cre Tg and IL-2 alleles, respectively. (M= Molecular weight ladder)

**Figure S4.**
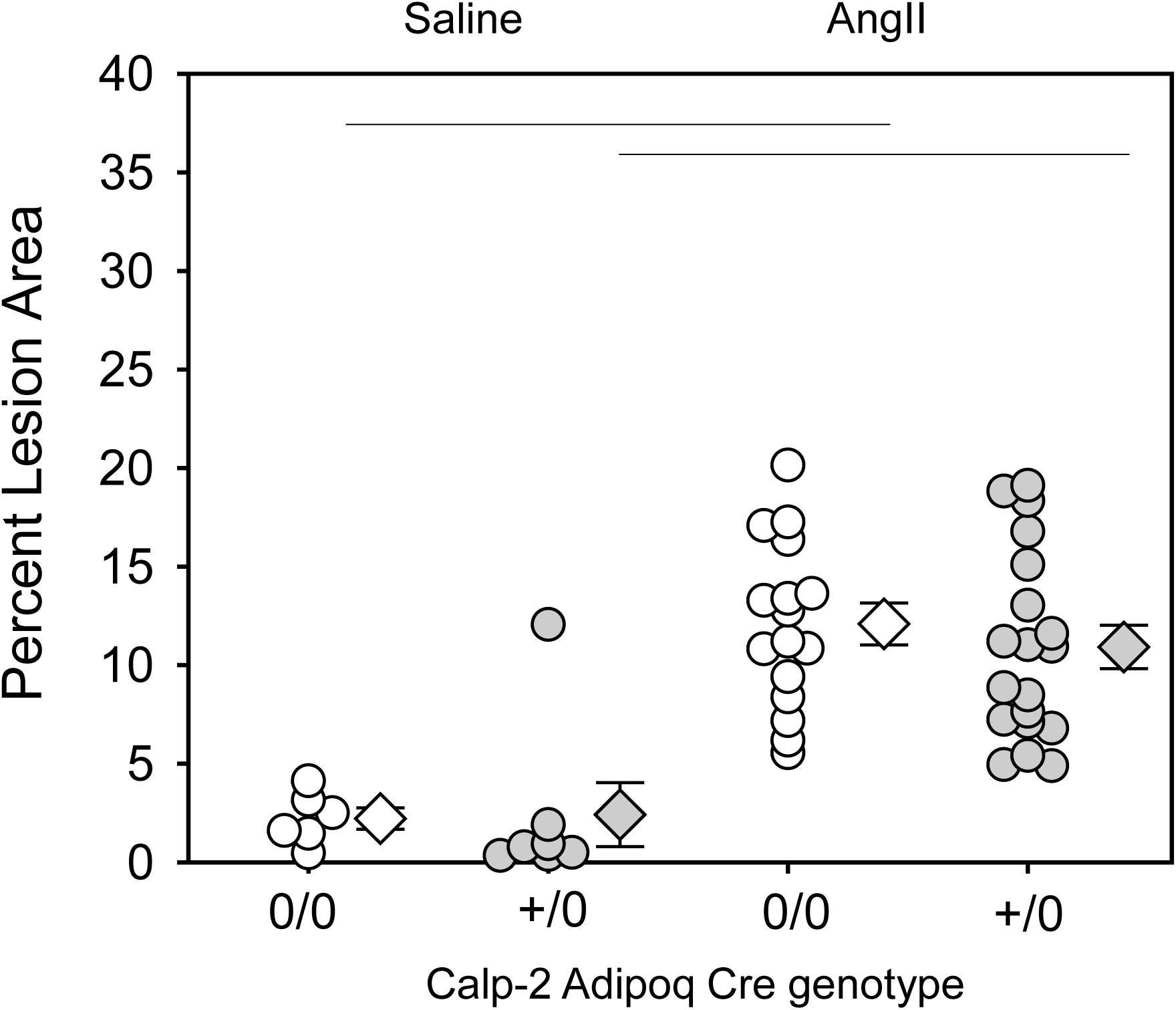
Calpain-2 deficiency in adipocytes had no effect on AngII-induced arch atherosclerosis. Atherosclerotic lesion area was measured on the intimal surface of aortic arch (**A**, Saline n=5; AngII n=16-20. White (Cre 0/0) and gray circles (Cre+/0) represent individual mice, diamonds represent means, and bars are SEMs. Horizontal lines represent significance of *P*<0.05 (Two-way ANOVA with Holm-Sidak post hoc analysis).

**Figure S5.**
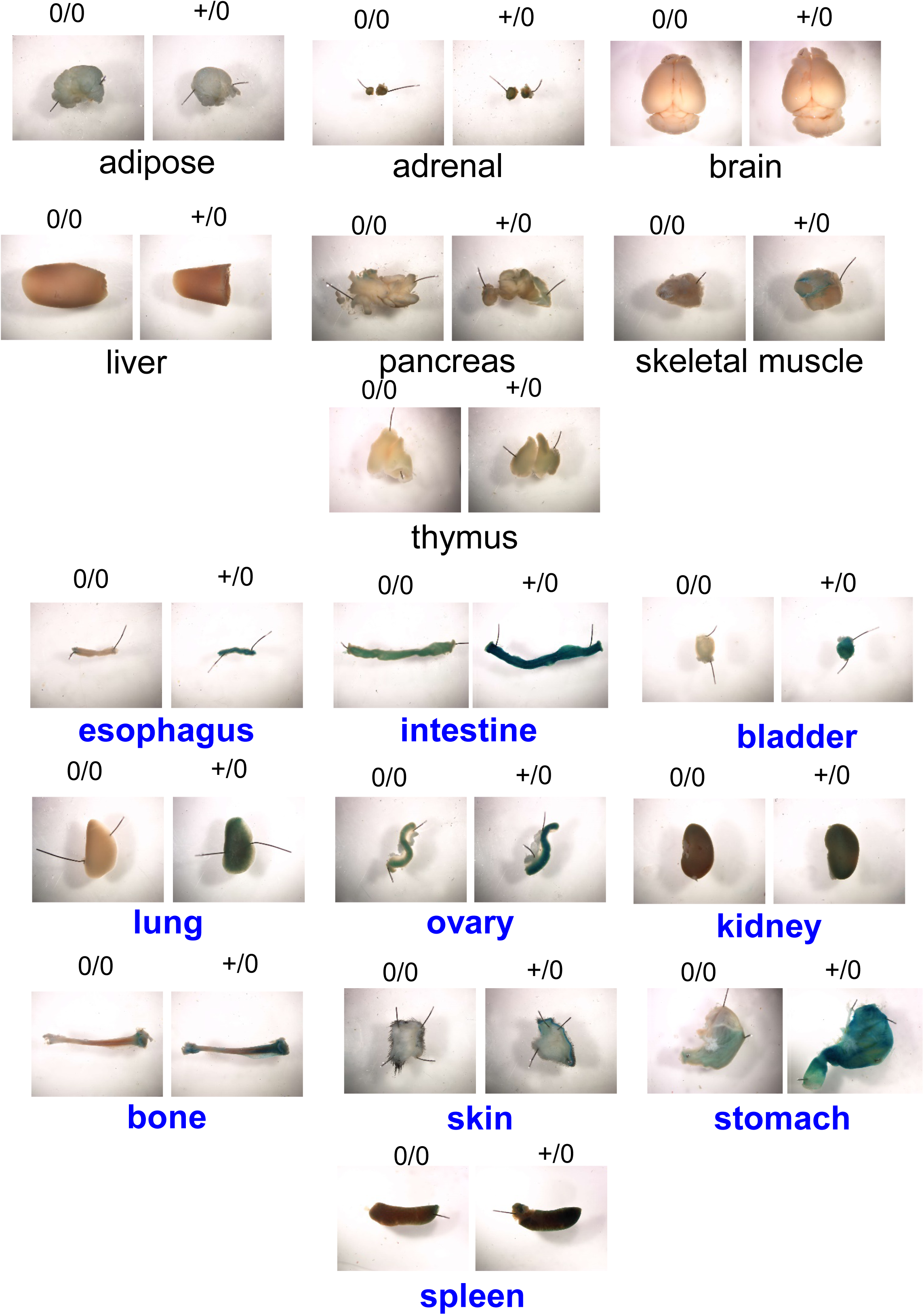
Representative X-gal staining images of various major organs from Col1a2 Cre 0/0 and Cre+/0 ROSA26R LacZ mice.

**Figure S6.**
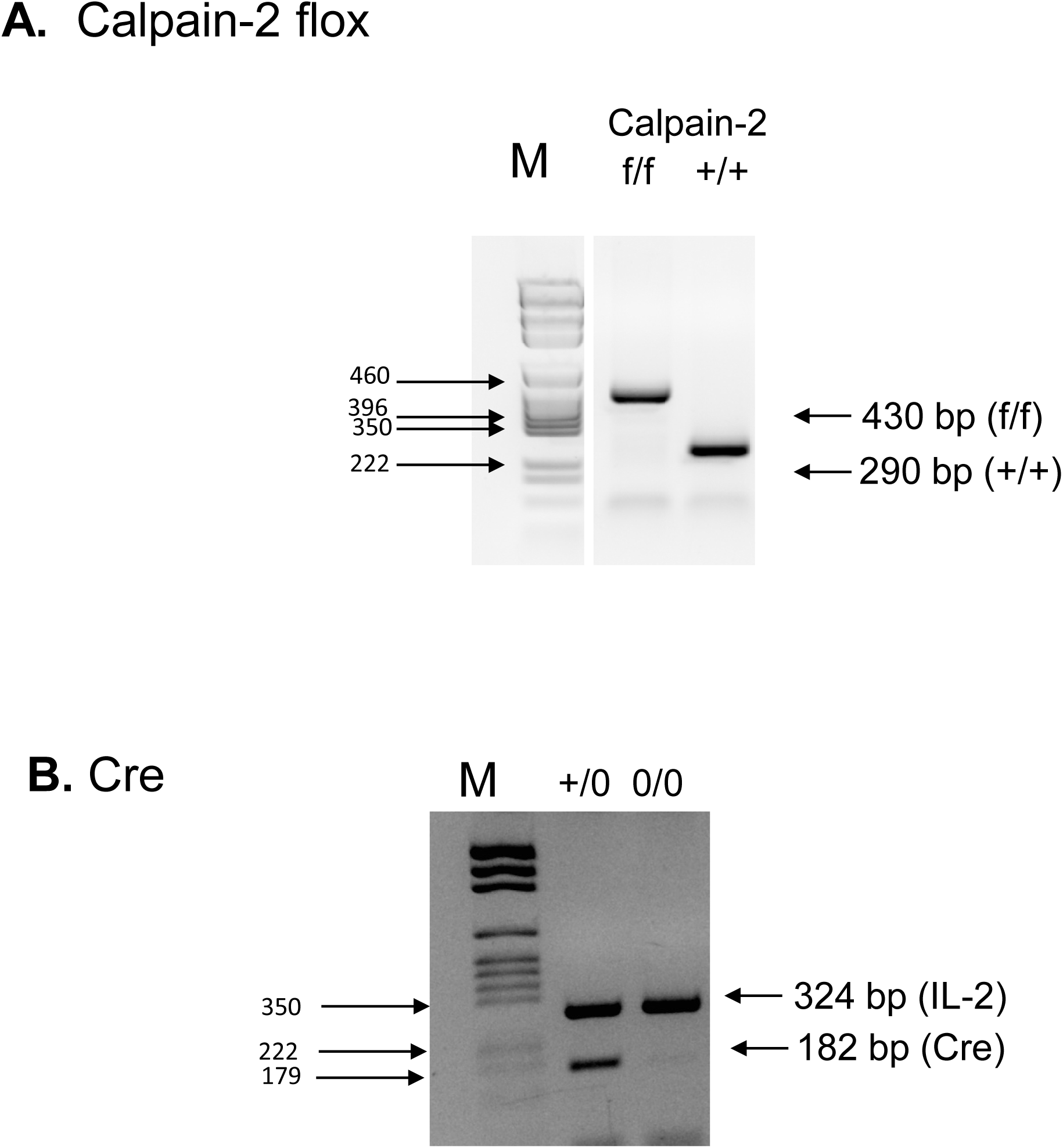
Genotyping of experimental mice for the calpain-2 floxed allele and Col1a2 Cre Tg by PCR. Genomic DNA from tail biopsies was isolated and screened by PCR. **A**. A 430 bp product was generated from calpain-2 f/f mice, while a 290 bp product was obtained from calpain-2 WT mice. **B**. Cre Tg was confirmed by PCR using IL-2 gene as internal control. PCR on genomic DNA yielded amplicons of 182 and 324 bp for Cre Tg and IL-2 alleles, respectively. (M= Molecular weight ladder)

**Figure S7.**
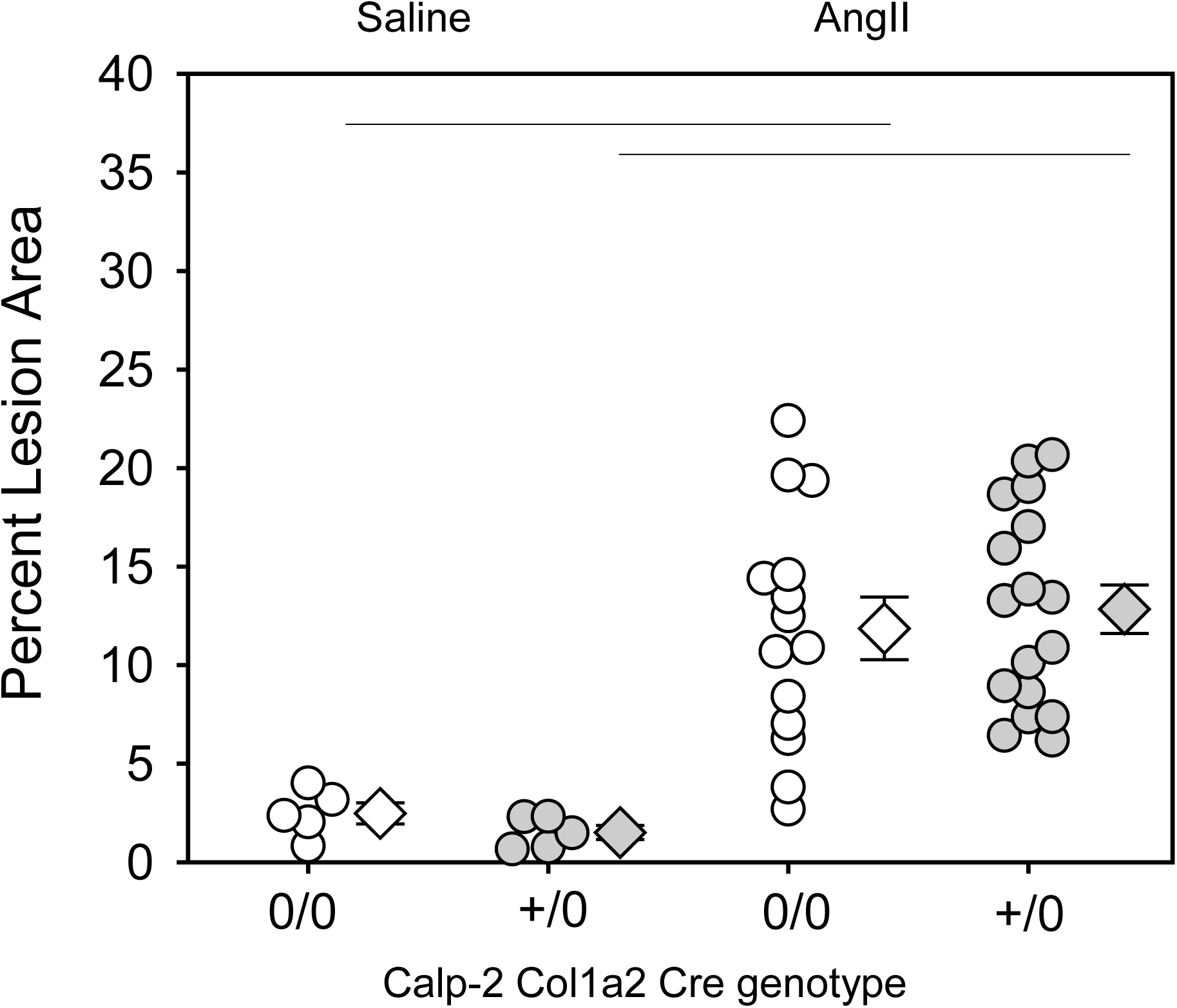
Calpain-2 deficiency in Col1a2 positive mesenchymal cells had no effect on AngII-induced arch atherosclerosis. Atherosclerotic lesion area was measured on the intimal surface of aortic arch (**A**, Saline n=5; AngII n=20-23. White (Cre 0/0) and gray circles (Cre+/0) represent individual mice, diamonds represent means, and bars are SEMs. Horizontal lines represent significance of *P*<0.05 (Two-way ANOVA with Holm-Sidak post hoc analysis).

**Figure S8.**
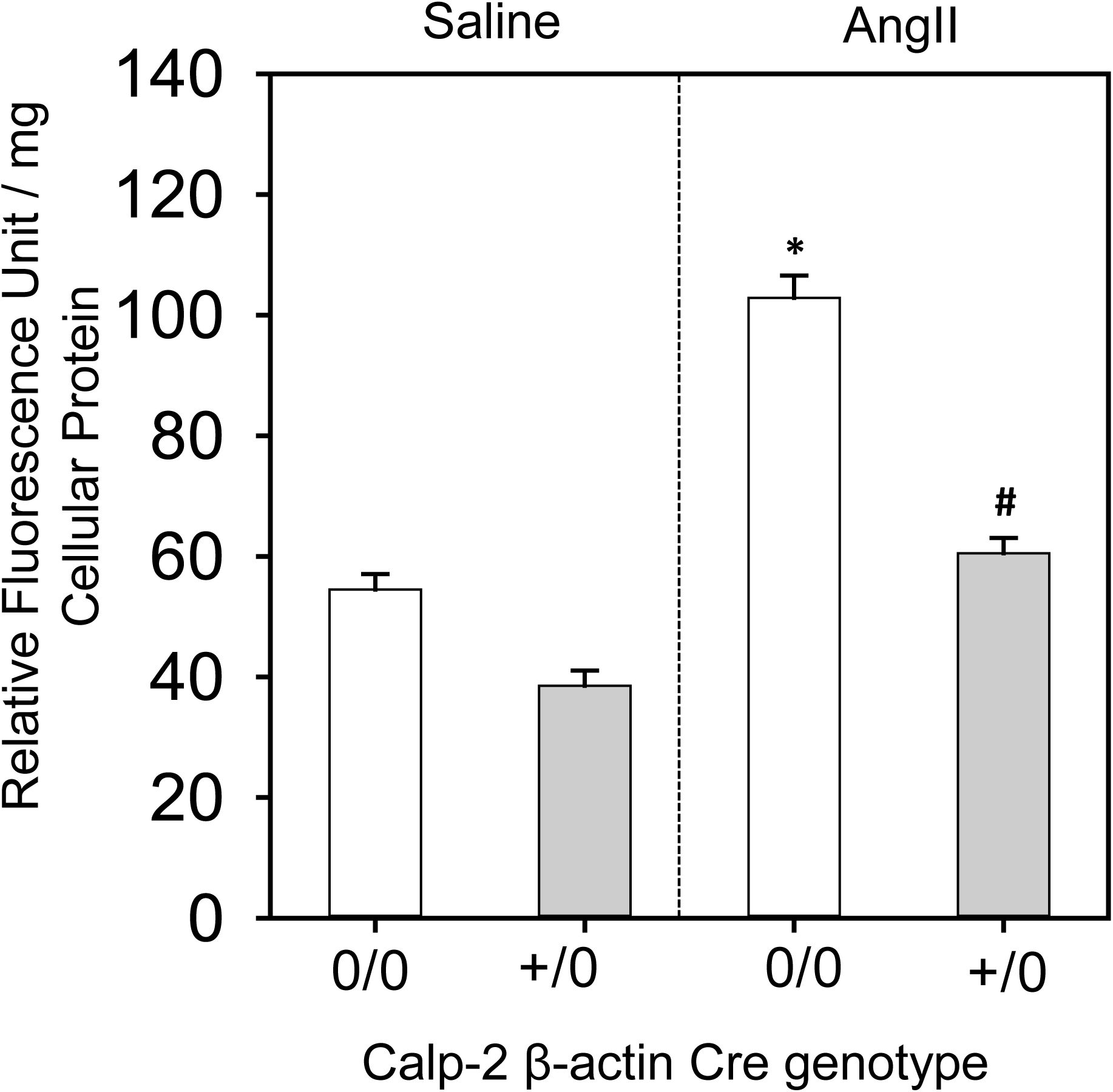
Calpain-2 deficiency suppressed AngII-induced calpain activity in the abdominal aorta. Calpain activity in abdominal aortas from saline and AngII-infused calpain-2 f/f β-actin promoter Cre0/0 and +/0 LDL receptor-/- mice (n=6/group). * denotes *P*<0.05 when comparing saline vs AngII infusion; # denotes *P*<0.05 when comparing Cre 0/0 vs Cre +/0 mice (Two-way ANOVA with Holm-Sidak post hoc analysis).

**Figure S9.**
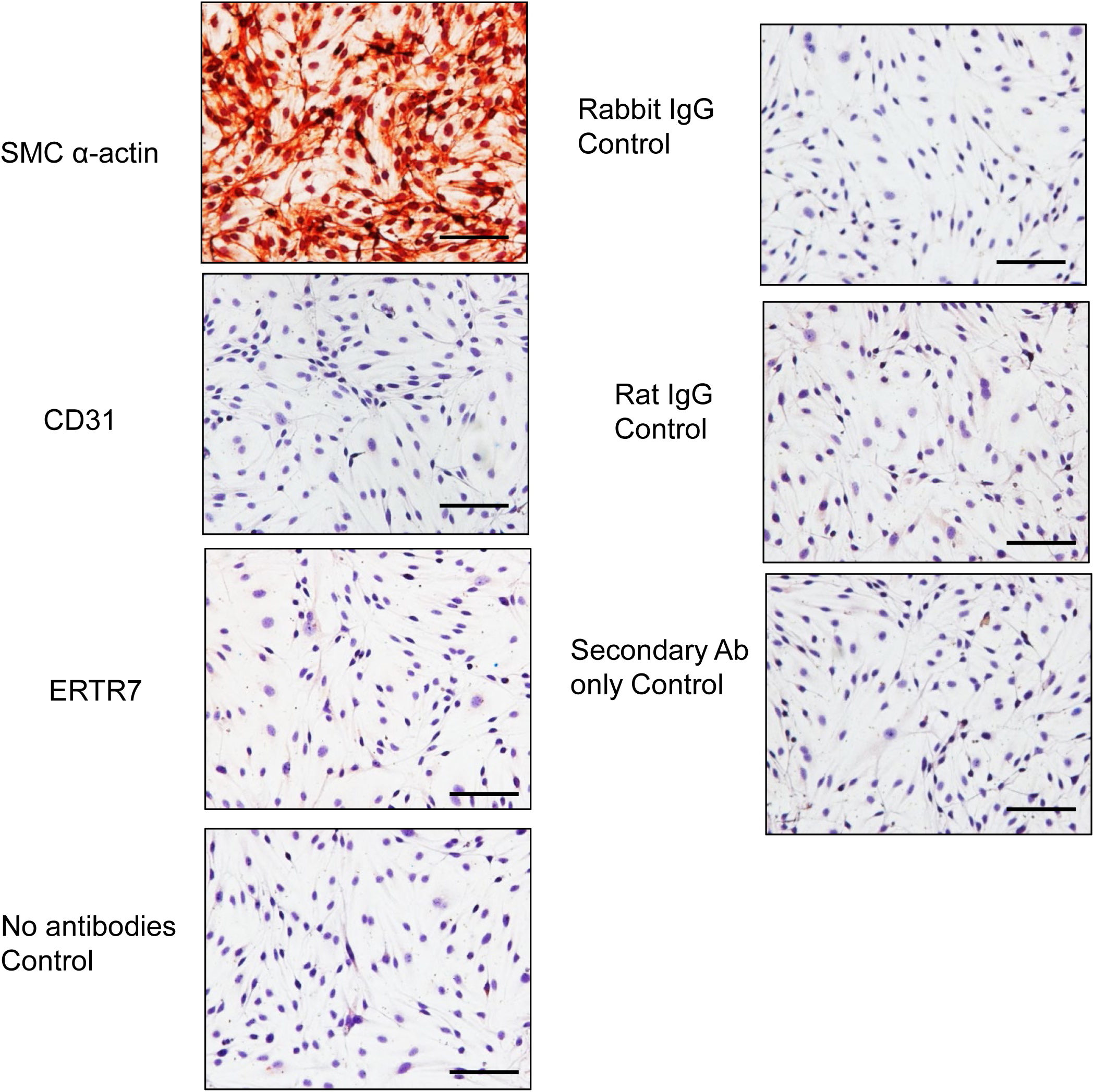
SMC α-actin staining on human aortic smooth muscle cells. Representative images of aortic SMCs immunostained for SMC α-actin, CD31 (endothelial cells), ERTR7 (fibroblasts) and iso—type control primary IgG, secondary antibodies controls. Positive cells stain red. Scale bars correspond to 50 μm (200x magnification).

**Figure S10.**
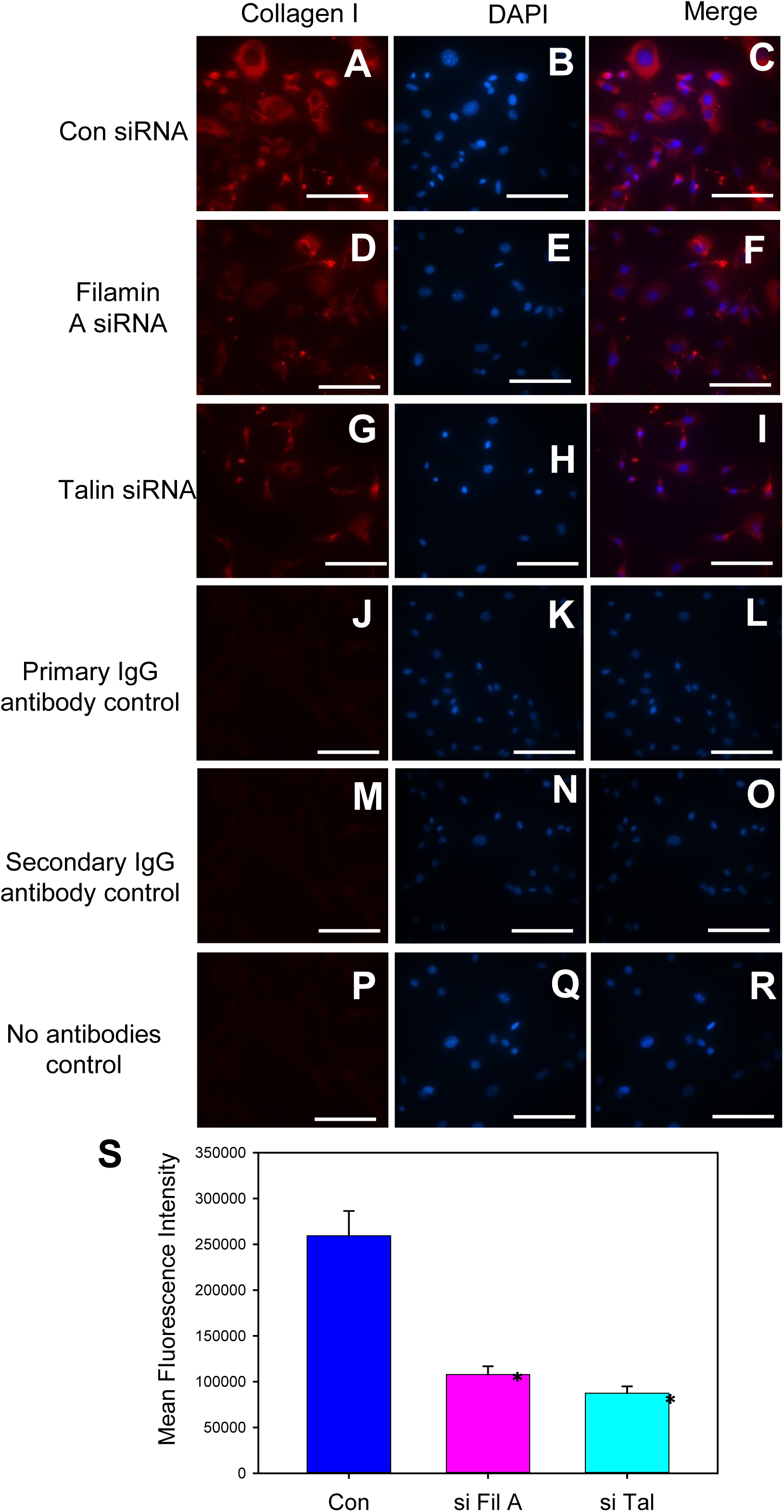
Silencing of filamin A or talin decreased collagen I in aortic SMCs. Immunofluorescent visualization of collagen I in control (**A-C**), filamin A (**D-F**) or talin (**G-I**) siRNA transfected mouse aortic SMCs. Representative SMC images stained with rabbit primary IgG iso-type control antibodies (**J-L**), secondary antibodies (Goat anti-rabbit-**M-O**) and no antibodies (**P-R**) control. Nuclei were counterstained with DAPI. Scale bars correspond to 50 μm (200x magnification). **S**. Quantification of mean fluorescence intensity. * denotes *P*<0.05 when comparing control vs filamin A or talin siRNA. (One-Way ANOV).A

**Figure S11.**
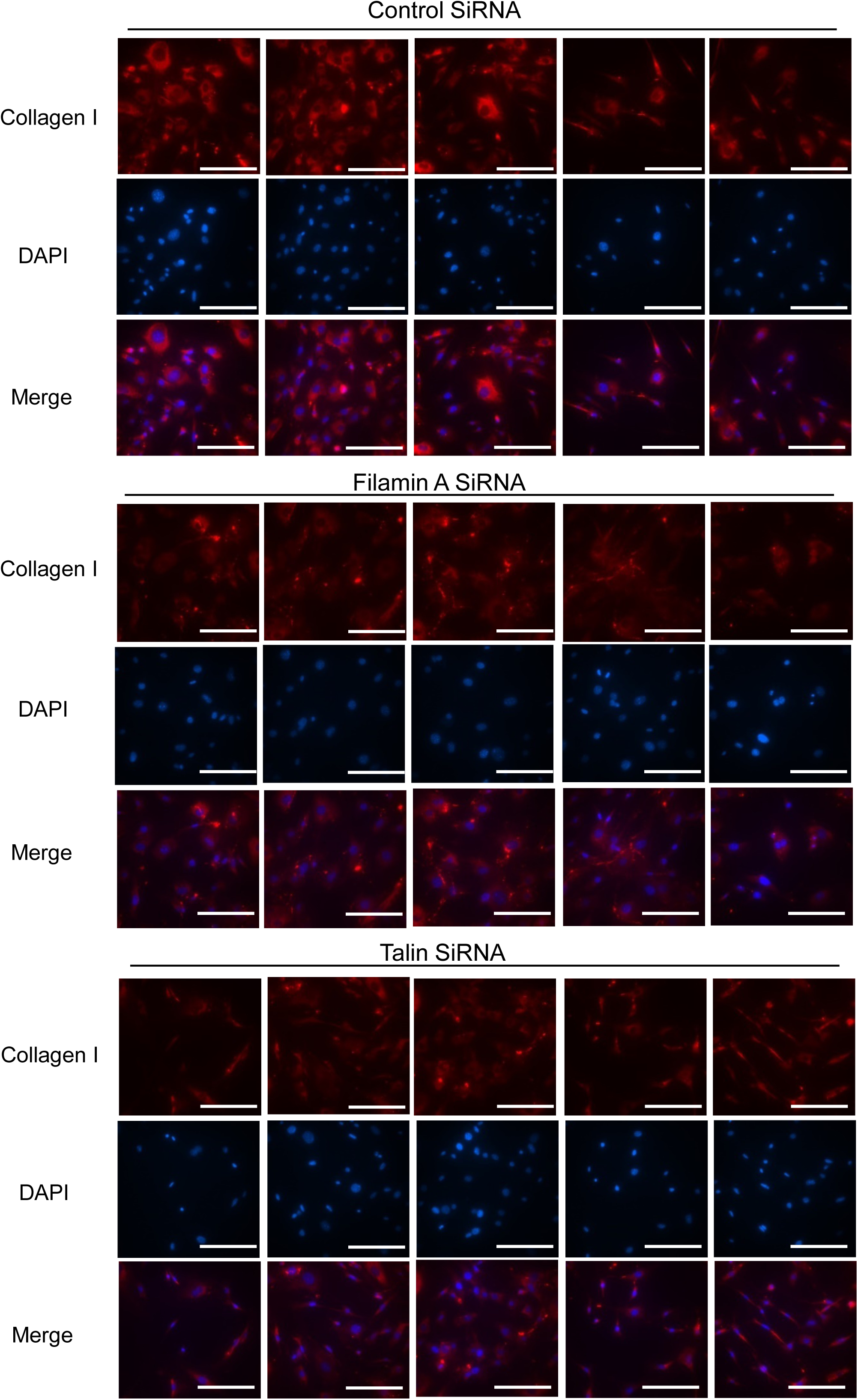
Representative images of collagen I staining in control, filamin A or talin silenced aortic SMCs.

## Notes

### Competing Interest Statement

The authors have declared no competing interest.

